# Differential transcriptomic response of *Anopheles arabiensis* to *Plasmodium vivax* and *Plasmodium falciparum* infection

**DOI:** 10.1101/2021.05.28.446219

**Authors:** Majoline Tchioffo Tsapi, Etienne Kornobis, Nicolas Puchot, Solomon English, Caroline Proux, Jessy Goupeyou-Youmsi, Anavaj Sakuntabhai, Marie-Agnes-Dillies, Randrianarivelojosia Milijaona, Romain Girod, Mamadou Ousmane Ndiath, Catherine Bourgouin

## Abstract

*Plasmodium vivax* malaria is now recognized as the second most dangerous parasitic threat to human health with the regular decrease of *Plasmodium falciparum* worldwide over recent decades. A very limited numbers of studies address the interaction of *P. vivax* with its *Anopheles* mosquito vectors. Those studies were conducted in *P. vivax* endemic countries with *P.vivax* local major vectors for which limited genomic and genetic tools are available. Despite the presence of *P. vivax* in several African countries and increasing reports on its occurrence in many others, there is virtually no data on the molecular responses of *Anopheles arabiensis,* a major African mosquito vector, to *P. vivax*, which limits the development of further “mosquito-targeted” interventions aimed at reducing *P. vivax* transmission. Taking advantage of the situation of Madagascar where *P. falciparum*, *P. vivax* and *An. arabiensis* are present, we explore the molecular responses of *An. arabiensis* towards these two human malaria parasites. RNA sequencing on RNAs isolated from mosquito midguts dissected at the early stage of infection (24 hours) was performed using mosquitoes fed on the blood of *P. vivax* and *P. falciparum* gametocyte carriers in a field station. From a *de novo* assembly of *An. arabiensis* midgut total RNA transcriptome, the comparative analysis revealed that a greater number of genes were differentially expressed in the mosquito midgut in response to *P. vivax* (209) than to *P. falciparum* (81). Among these, 15 common genes were identified to be significantly expressed in mosquito midgut 24 hours after ingesting *P. vivax* and *P. falciparum* gametocytes, including immune responsive genes and genes involved in amino-acid detoxification pathways. Importantly, working with both wild mosquitoes and field circulating parasites, our analysis revealed a strong mosquito genotype by parasite genotype interaction. Our study also identified 51 putative long non-coding RNAs differentially expressed in *An. arabiensis* mosquito infected midgut. Among these, several mapped to the published *An. arabiensis* genome at genes coding immune responsive genes such as gambicin 1, leucine-rich repeat containing genes, either on sense or antisense strands.

This study constitutes the first comparison of *An. arabiensis* molecular interaction with *P. vivax* and *P. falciparum*, investigating both coding and long non-coding RNAs for the identification of potential transcripts, that could lead to the development of novel approaches to simultaneously block the transmission of *vivax* and *falciparum* malaria.

## Introduction

Malaria is still a major public health concern in many tropical and subtropical countries (WHO 2020). Although malaria eradication has emerged as a goal for the next decade, the field consensus is that the development of novel intervention strategies is hindered by our still limited understanding of the biology of *Plasmodium*, the causative agent of malaria, and of the complex interactions that the parasite maintains with its mammalian and mosquito hosts.

With more than 2-decade of intense focus on *Plasmodium falciparum*, which is responsible for the majority of deaths, major progresses have been achieved leading to substantial *falciparum* malaria decrease, mostly outside Africa (WHO 2020). This happened despite the worrying appearance of artemisinin-resistant *P. falciparum* parasites in South-east Asia (Menard and Dondorp 2017). *Plasmodium vivax* is the most geographically widespread species among the human malaria parasites infecting an 14-25 million people annually (Howes, Battle, et al. 2016; Battle et al. 2019). Commonly considered as a mild disease, *P. vivax* malaria is responsible for numerous severe cases, the frequency of which is now clearly reported (Anstey et al. 2012; Baird 2013; Rahimi et al. 2014; Bourgard et al. 2018; Mukhtar et al. 2019).

Occurrence of *P. vivax* malaria is overall rare in Sub-Saharan Africa submerged by the deadly *P. falciparum* malaria (Battle et al. 2019; Weiss et al. 2019). It has classically been considered that *P. vivax* parasites were unable to infect individuals from black ancestry and further argued that absence of *P. vivax* in population from black ancestry was linked to the absence of the *P. vivax* receptor (the Duffy Antigen Receptor for Chemokines, DARC) on the red blood cells of those individual (Miller et al. 1978). In the few African countries with admixed populations of diverse genetic ancestry (Mauritania, Ethiopia and Madagascar), *P. vivax* is often the second cause of malaria (Solomon, Kahase, and Alemayehu 2020; Howes, Mioramalala, et al. 2016; Ba et al. 2016). In those countries, the same mosquito species do transmit both parasite species, with *Anopheles arabiensis* and its sister taxa *Anopheles gambiae* as major vectors mostly known as the major vectors of *P. falciparum* in sub-Saharan Africa.

Facing the absence of an efficient vaccine, increased resistance of malaria parasites to artemisinin-based combined therapies (ACT) and increased resistance of mosquitoes to insecticides, transmission-blocking approaches have been placed at the forefront of additional strategies to combat malaria (Rabinovich et al. 2017). Transmission blocking (TB) strategies come in two “flavours”: targeting the parasite stages developing into mosquitoes or targeting mosquito molecules essential for malaria parasite development into their vectors (Lavazec and Bourgouin 2008; Saul 2007). Recent advances in mosquito transgenesis mixing both parasite and mosquito targets have demonstrated that combined approaches might be feasible (Isaacs et al. 2012). Transmission blocking approaches based on mosquito molecules have almost exclusively focused on *P. falciparum* in the last decades with numerous studies of *P. falciparum* interaction with its major African vector *An. gambiae,* for which molecular and genetic tools have been developed. Today a limited number of targets has been proposed as target for transmission blocking strategies. Most of these targets correspond to mosquito molecules expressed in the mosquito midgut, the compartment where malaria parasites initiate their sporogonic development (Dinglasan et al. 2007; Lavazec et al. 2007; Zhang et al. 2015). Indeed, during the parasite early sporogonic development a limited number of ookinetes, the midgut invasive stage, are produced, representing a weak point for efficient intervention (Gouagna et al. 1998; Smith, Vega-Rodríguez, and Jacobs-Lorena 2014). Target validation is usually accomplished by experimental mosquito feeding on *Plasmodium-* infected blood supplemented with antibodies raised against the candidate transmission- blocking molecule. As *P. vivax* is not cultivable yet, validation of TB candidates derived from *An. gambiae-P. falciparum* studies were performed in *P. vivax* endemic countries and its local vectors. These resulted in contrasting results of modest robustness due to the regulatory difficulty to include *An. gambiae* -*P. falciparum* as an internal control. Nevertheless, ingestion of antibodies against *An. gambiae* APN1 and FRET1 reveals similar trends in reducing *P. falciparum* and *P. vivax* in *An. gambiae* and *Anopheles dirus*, respectively (Armistead et al. 2014; Niu et al. 2017). On the contrary, antibodies against AgSGU, an *An. gambiae* midgut GPI-anchored protein reduced development of *P. falciparum* in *An. dirus* but had no effect on *P. vivax* development in this vector (Mathias et al. 2014). This latter data is in line with previous ones that suggest that the development of *P. falciparum* and *P. vivax* in *Anopheles tessellatus* involves both common and different molecular mechanisms (Ramasamy et al. 1996).

With the advances of Next Generation Sequencing (NGS) technologies, transcriptomic approaches are being developed to gain further insights into *Anopheles* responses to *P. vivax* (Santana et al. 2019; Boonkaew et al. 2020; Kumari et al. 2021). Although providing substantial novel data, these studies were conducted with local *Anopheles* vectors for which limited molecular information or genetic tools are available and did not address the specificity of the mosquito response to either human *Plasmodium* species. Taking advantage of the situation of Madagascar where both *P. falciparum* and *P. vivax* are responsible for malaria and are transmitted mostly by *An. arabiensis* we performed a differential midgut transcriptomic analysis of local *An. arabiensis* mosquitoes infected with circulating strains of *P. falciparum* and *P. vivax* with the aim to identify molecules or pathways that could constitute targets for transmission-blocking approaches of either or both *P. vivax* and *P. falciparum*. At the initiation of the project, an RNA-Seq pilot study revealed a substantial sequence divergence of the Malagasy *An. arabiensis* RNA complement with the published *An. arabiensis* annotated genome that used a mosquito colony established from Sudan origin (VectorBase). Therefore, we chose to perform a *de novo* transcriptome assembly from our complete sets of RNA-Seq data. Our differential analysis reveals that 209 genes were differentially expressed in *An. arabiensis* midgut cells in response to *P. vivax* whereas only 81 were found differentially expressed upon *P. falciparum* infection, among which several transcripts encoding lncRNAs. A limited number of genes were regulated by both parasite species, which might facilitate selection of targets for limiting *Plasmodium* transmission.

## Materials and Methods

### Study area and mosquito production

The study took place in a field station established in Andriba (17°35’49.92” S, 46°56’0.59” E), Maevatanana district in the Northwest of Madagascar (Goupeyou-Youmsi et al. 2020). In this area, malaria is endemic and mainly due to *P. falciparum* and *P. vivax*, with *P. falciparum* being the most prevalent parasite species. The highest intensity of transmission is observed from November to April (Nguyen et al. 2020). Major and secondary Malagasy malaria vectors are present in the area: *An. gambiae*, *An. arabiensis*, *Anopheles funestus*, *Anopheles mascarensis* and *Anopheles coustani*. Anopheline larvae and pupae were collected daily in rice fields and water ponds in the Andriba village, using the standard dipping technique (Service 1993). After removal of non-anopheline larvae and predators, larvae and pupae were placed in clean water and larvae fed on Tetramin Baby and grounded cat-food until adult emergence. Morphological identification of adult mosquitoes revealed that over 95% of the collected larvae and pupae were indeed *A. gambiae s.l.* which includes both *An. gambiae s.s and An. arabiensis,* two morphologically indistinguishable sibling species. Only those adults were maintained in 30×30 cm cages with *ad libidum* access to a 10% sucrose solution, in the field insectary. Day-light conditions of the insectary matched the natural local day-night cycle close to 12:12. Humidity was controlled by regular floor watering. In addition, adult cages were covered with wet tissue and placed behind plastic curtains to limit dryness and too high temperatures. Day-time temperature in the insectary matched roughly the local temperature while night-time temperature decreased by a few degrees only, by keeping the insectary doors tightly closed. Both temperature and humidity were recorded daily. Overall, temperature ranged from 24 to 28 °C and humidity close to 80%.

### Recruitment of *Plasmodium vivax* and *Plasmodium falciparum* gametocyte carriers

Recruitment of *P. vivax* gametocyte carriers was performed among symptomatic patients attending the health centers of Antanimbary (17°11’06.6“S 46°51’19.4”E) and Andriba (17°35’49.92” S, 46°56’0.59” E). Indeed, *P. vivax* gametocytes develop at any red-blood cell invasion cycle and are therefore, found in symptomatic individuals. On the contrary, *P. falciparum* mature gametocytes need 10 to 15 days to develop as an escape strategy away from the initial disease symptoms. Therefore, *P. falciparum* gametocyte carriers were selected among asymptomatic 5- to 14-year-old children who attended the local primary schools. For each child, a blood sample obtained by finger-prick was used to perform a RDT (malaria Pf/pan SD Bioline), thick and thin blood smears. The RDTs allow detection of *P. falciparum* and any of the others *Plasmodium* parasites, *P. vivax*, *Plasmodium ovale* and *Plasmodium malariae,* which all are present in Madagascar. Microscopy analysis was used to determine *Plasmodium* species and parasites stages. Asexual (trophozoïtes) and sexual (gametocytes) stages of *P. vivax and P. falciparum* were detected by light microscopy (100X magnification) on 10% Giemsa-stained thick blood smears. *Plasmodium* gametocyte density was determined against 500 white blood cells (WBC), assuming the standard number of 8,000 WBC/μl of blood, using the 10% Giemsa-stained thin blood smears. An Artemisinin-based combination therapy (ACT) was given to each child with a positive RDT or carrying *Plasmodium* trophozoïtes, as evidenced by microscopy, according to the national guidelines. Children identified as *P. vivax* or *P. falciparum* gametocyte carriers were enrolled for providing blood for mosquito infections after their parents or legal guardians had signed documents indicating informed consent. All procedures involving human subjects used in this study were approved by the national ethical committee of the Malagasy Ministry of Health (N° 122-MSANT/CE).

### Experimental design

Our global objective was to compare the mosquito midgut response to *P. vivax* and to *P. falciparum*. Because we worked with circulating *Plasmodium* parasites, *Plasmodium* isolates used for replicate infection experiments are likely different from each other. To be able to study mosquito midgut transcriptome regulation in response to the developing parasites, our experimental strategy requires performing two concurrent mosquito infections for each gametocyte sample (Figure S1). The first infection was performed with “infective” gametocytes and the second with heat inactivated gametocytes, there after termed “non-infective” gametocytes. In addition, as we worked with F0 from wild mosquitoes that likely have different genetic backgrounds, we aimed to consider individual midgut samples as mosquito biological replicates per gametocyte sample. Further, to avoid technical bias, individual mosquito midguts were selected from 2 (*P. vivax*) and 3 (*P. falciparum*) technical replicates per gametocyte sample. Overall, we selected 4 independent infections for each parasite species that resulted in prevalence of infection close to or above 40% as determined by oocyst detection in the “infective sample” 7 days post infection. After RNAs isolation and quality control 16 individual midguts for *P. vivax* and 24 individual midguts for *P. falciparum* were processed for RNA-Seq.

### Mosquito experimental infections

Batches of 50 to 70 females 3 to 5 days-old were dispatched in homemade feeding pots and starved from sugar 16 hours before infection. From selected *Plasmodium* gametocyte carriers 5 ml of venous blood were drawn in heparinized tubes. The blood sample was centrifuged at 2,000 rpm, 3 min and 37°C and the patient serum was replaced with AB serum from donors never exposed to malaria (EFS-France) to limit possible effect of acquired transmission-blocking immunity (Boudin et al. 2005). The blood sample was divided into two parts. The first part was directly proposed to mosquitoes and constitutes the “infective gametocyte sample”; the second part was heated and shaken at 42 °C for 20 minutes (Eppendorf Thermomixer Comfort) to inactivate *Plasmodium* gametocytes before being proposed to mosquitoes. This constitutes the “non-infective gametocyte sample” (Figure S1). Heat treatment was previously demonstrated to fully inhibit gametocyte infectivity (Mendes et al. 2008).

Membrane feeding assay was carried out essentially as previously described (Tchuinkam et al. 1993), using stretched Parafilm^®^ and glass feeders maintained at 37°C. Female mosquitoes were allowed to feed for 60 min, in the dark. Fully fed females were transferred into small cages, matching each experimental pot, and given free access to a 10% sucrose solution. At 24 hours post blood meal (PBM), individual midguts were dissected in cold 1X PBS, transferred into 1.5 ml micro-tube containing 100μl of Tri-Reagent (MRC) and stored at −20°C for few days before being stored at −80°C at the Institut Pasteur de Madagascar. Carcasses from each dissected mosquito were individually stored in micro-tubes for further species identification. The remaining mosquitoes were maintained under optimal conditions in the insectary and their midgut dissected on day 7 PBM to determine prevalence and load of infection by counting oocysts on mercurochrome (0.4% w/v- Sigma-Aldrich-M7011) stained midguts using a light microscope (X20 & X40).

At the end of the transmission season, all biological samples were sent under dry ice to Institut Pasteur in Paris for RNA-Seq analysis.

### Mosquito species identification

Although we had previous indication that the majority of the *A. gambiae s.l.* present in Andriba belongs to the *An*. *arabiensis* taxon, we used a PCR based procedure to confirm that indeed all infected samples were from *An. arabiensis* mosquitoes. For that, DNA was extracted from each mosquito carcass matching the prepared midgut samples, using DNAzol**®** reagent (MRC). PCR primers were as previously published (Fanello, Santolamazza, and della Torre 2002) (Table S3). Mosquitoes from our *An. gambiae* Yaoundé colony and a recently established *An. arabiensis* colony were used as control.

### RNA extraction and reverse transcription

Individual mosquito midguts stored in 100μl Tri-Reagent were crushed using polypropylene micro-pestles (Dutscher 045007). Total RNA purification was performed using the Direct-zol™ RNA Microprep isolation kit (Zymo Research, Cat # R2062), according to the manufacturer’s instructions. The Tri-Reagent extract was loaded onto Microprep column and total RNA was eluted with 10μl of DNase/RNAse free water and further treated with DNAse (Invitrogen™ TURBO DNA-free™ Kit). RNA quantification was performed using Nanodrop One (Thermo Scientific).

Before initiating the RNA-Seq experiments, we verified that every single “infected” midgut contained developing *Plasmodium* parasites by detecting the ookinete mRNA marker encoding Pvs25 or Pfs25 using primer sets reported by (Wampfler et al. 2013) and q-PCR (Power SYBR™ Green PCR Master Mix amplification - Life Technologies).

Reverse transcription reactions were performed in 40μl final volume using 3 μl of RNA, 8μl of 5X First-Strand Buffer, 4μl 0.1M DTT, 1μl of 20mM dNTPs, 2μl of 50 μM random hexamers, 1μl of 40U/μl RNaseOUT™ recombinant Ribonuclease Inhibitor and 2μl of 200 Units MMLV Reverse Transcriptase. All reagents were from Life Technologies Invitrogen, except Random hexamers obtained from Promega.

### RNA-Seq library preparation and Illumina sequencing

RNA quality was assessed using the Agilent Bioanalyzer. Briefly, selected RNA samples were denatured, by incubating at 70°C for 2 minutes, and then placed on ice. Nine μl of the gel-dye mix was pipetted into the bottom of a nanochip. Then, 1μl of samples, 1μl of RNA 6000 Nano Marker and 1μl of RNA 6000 Ladder were pipetted into assigned well, vortexed and run using 2100 expert Eukaryote Total RNA Nano Series II (Agilent Technologies).

Forty RNA samples of good quality obtained from infected (infective gametocytes) and cognate uninfected (heat-inactivated gametocytes) individual mosquito midguts were selected for cDNA library preparation: 16 corresponding to *P*. *vivax* and 24 to *P. falciparum* (Figure S1). RNA-Seq library preparation was performed using the SMARTer^®^ Stranded Total RNA-Seq kit – Pico Input Mammalian (Takara Bio USA, Inc) with the RNA input ranging from 7 to 10 ng of starting material.

The protocol starts with first-strand cDNA synthesis from total RNAs fragmented for 3 minutes following Takara’s recommendation. The Illumina adapters and indexes were added, followed by the purification of the RNA-Seq library and depletion of ribosomal cDNA with ZapR and R-Probes. The final RNA-Seq library was obtained after 15 cycles of PCR amplification and purified using Agencourt AMPure XP beads (Beckman and Coulter, Beverly, MA, USA). The library size distribution was evaluated by running samples on the Agilent 2100 Bioanalyzer. The 40 libraries were then sequenced using the HiSeq 2500 (Illumina, San Diego, CA, USA) at 2×100 bp paired-end sequencing with v4 chemistry. The 40 samples were distributed on the 8 lanes of the flow cell such that there was no confounding effect between the lanes and the biological factors of interest. Library preparation and sequencing were performed at the Institut Pasteur Epigenetics and Transcriptomics Platform (PF2).

### *de novo* assembly of *An. arabiensis* midgut transcriptome

From a pilot study using STAR 2.7.0d and the available assembly of the *An. arabiensis* genome on VectorBase, only 25% of the RNA-Seq reads from our Malagasy *An. arabiensis* samples could be mapped. In addition, functional annotation was scarce. Therefore, we performed a *de novo* midgut transcriptome assembly using the paired-end sequencing.

Quality control benefited from MultiQC 1.7 (Ewels et al. 2016). The resulting reads (30–90 million) displayed in Table S2 were quality trimmed and adapters clipped using fastp 0.19.6 (Chen et al. 2018) prior to *de novo* assembly using Trinity 2.6.6 (Grabherr et al. 2011). Completeness of the transcriptome was evaluated by BUSCO 3.0.2 (Simão et al. 2015), comparing with *Diptera* and *Insecta* reference transcriptomes.

### Functional annotation and differential expression analysis

Functional annotation of the midgut transcriptome was performed with Trinotate 3.1.1 (Bryant et al. 2017) which uses Blast 2.5.0 (Camacho et al. 2009), HMMER 3.2 (Finn, Clements, and Eddy 2011), TMHMM 2.0 (Krogh et al. 2001), SignalP 4.0 (Petersen et al. 2011) and RNAmmer 1.2 (Lagesen et al. 2007). Two groups of transcribed features were obtained: annotated features originating from known coding genes and unannotated features that could correspond to previously undescribed genes or non-coding RNA. The PLEK 1.2 (Li, Zhang, and Zhou 2014) and CPAT 1.2.4 (Wang et al. 2013) non-coding RNA identification programs were next used to refine the unannotated features. In order to investigate further their function, putative lncRNA transcripts were mapped to *An. arabiensis* Dongola AaraD1 genome and its annotation AaraD1.11 to identify potential target genes.

Transcript expression levels comparing infected versus uninfected *An. arabiensis* mosquitoes were quantified using kallisto 0.45.0 (Bray et al. 2016). Transcript abundances were used for gene-level comparisons using tximport 1.6.0 (Soneson, Love, and Robinson 2015) from the three and four biological replicates of *P. vivax* infection and *P. falciparum* infection. DESeq2 1.18.1 (Love, Huber, and Anders 2014) was used to detect genes differentially expressed between the infected and uninfected mosquito midgut samples. Differentially expressed genes were further analyzed for functional classification using Gene Ontology (GO). GO enrichment analysis was performed using Trinity scripts and GOSeq 1.30.0 (Young et al. 2010) to provide a shortlist for potential genes of interest classified in three categories: biological process, molecular function and cellular component.

### Code and data availability

The bioinformatic analysis will be provide later as Jupyter notebooks. The RNA-Seq reads, metadata and raw reads count will be available after submission of the manuscript for publication. Differential expressed genes and GO term analysis for each *Plasmodium* isolate are also presented in Supplementary Tables 4 to 6.

## Results

### Selection of mosquito midgut samples for the differential RNA-Seq analysis

From 14 *An. arabiensis* experimental infections performed in our field station between January and March 2017, we selected 4 infections with *P. vivax* and 4 infections with *P. falciparum* (Table S1 and Figure S2). The inclusion criteria were enough individual dissected mosquito midguts 24h post blood meal, prevalence of infection at day 7 post infection in the experimental “infective” blood meal close to or above 40% and absence of parasites in the “non-infective” blood meal at day 7 post infection. Although *Pv3* exhibited a reduced prevalence of infection at d7 (17.5%) it was initially included in our analysis. The second criterion was detection of the mRNA encoding the *Plasmodium* ookinete surface protein 25 (*Pvs25* or *Pfs25*) in individual midguts by RT-qPCR. The last criterion was the quality of the RNA extracts for subsequent RNA-Seq library construction as determined using the Agilent Bioanalyzer. Overall, as depicted in Figure S1, 16 RNA-Seq libraries were produced from mosquitoes challenged with *P. vivax* and 24 from mosquitoes challenged with *P. falciparum.*

### Summary of RNA-Seq raw data

Sequencing of the 40 libraries resulted in a total of 1.3 billion paired-end reads. From the paired-end reads, *de novo* midgut *An. arabiensis* transcriptome assembly was carried out and produced 250642 transcripts; 112242 of them being supported by at least 5 reads and correspond to 164954 putative genes (i.e. trinity clusters). Completeness of the transcriptome was evaluated at 78% and 94% by BUSCO, respectively comparing with *Diptera* and *Insecta* reference transcriptomes. Annotation performed by the Trinotate suite led to 60243 transcripts showing significative similarity with swissProt or Pfam entries, 12083 having a blastx hit with *Drosophila melanogaster* nucleotide sequence. From the transcripts, we performed a Principal Component Analysis (PCA) based on VST (variance stabilizing transformation) transformed counts from DESeq2 to visualize the overall effect of the infected and the uninfected status of the mosquito midguts according to each *Plasmodium* species and isolate (Figure S3). We excluded from the PCA, *Pv3* samples as the corresponding libraries had much less sequencing coverage than the others. Considering that the mosquito samples originated from different field collections across the malaria transmission season and that the parasite isolates came from different gametocyte carriers, it was anticipated that significant genetic variation could be observed among mosquito samples and among parasite isolates. The PCA shows that components 1 and 2 together display 58 % and 52 % of the total variance in response to *P. vivax* and *P. falciparum* infection, respectively.

Three clusters were clearly identified representing the three independent *P. vivax* mosquito infections (Figure S3A). Nonetheless, no clear structure was observed corresponding to the infected/uninfected status of the mosquitoes. The results were slightly different with *P. falciparum* with a trend of clustering between infected and uninfected mosquitoes within a *Pf* isolate (Figure S3B). More striking was the clustering of *Pf1* and *Pf2* data. Indeed, *Pf1* and *Pf2* infections used the same batch of mosquitoes, suggesting a strong contribution of mosquito genetic component. Based on this PCA analysis, and since there was extensive variation within groups and no clear overall segregation between infected and uninfected mosquito midguts, we performed individual differential analyses using each parasite isolate separately (*Pv*1, *Pv*2, *Pv*4*, Pf*1, *Pf*2, *Pf*3 and *Pf*4) comparing infected versus uninfected mosquito samples.

### Differential gene expression in *An. arabiensis* midguts infected with *P. vivax*

Using a False Discovery Rate of 0.05 (FDR < 0.05) as the threshold to classify differentially expressed genes, our analysis resulted in the identification of 209 transcripts differentially regulated in *An. arabiensis* midguts 24h after feeding on infective *P. vivax* gametocytes compared to mosquito fed on cognate un-infective gametocytes. The detailed gene list is provided in Table S4. From the 209 transcripts identified in mosquito midguts as differentially expressed in response to *P. vivax* isolates, 36 were up-regulated and 173 genes were down-regulated (Figure 1A). The majority of the transcripts were specific to each *P. vivax* isolate. Five down-regulated transcripts were shared between *P. vivax* isolates 1 and 4 (Figure 1B and Table 1). The most down-regulated *An. arabiensis* gene was TRINITY_DN109598_c5_g1 with two transcripts showing a log_2_FC at - 6,357 identified in *Pv1* infection. This gene showed 36.89% identity to Aminopeptidase N_bovin. Another down regulated transcript, TRINITY_DN108067_c0_g1 in *Pv2* infection showed 98.04% identity to *An. gambiae* Defensin 1 with log_2_FC at - 4.13. The most up-regulated transcripts were TRINITY_DN101897_c5_g2 with three variants showing log_2_FC at 4.549 identified in *Pv4* infection, and TRINITY_DN103245_c1_g1 showing log_2_FC at 4.551 in *Pv1* infection; both transcripts were classified as unannotated genes.

**Table 1.**
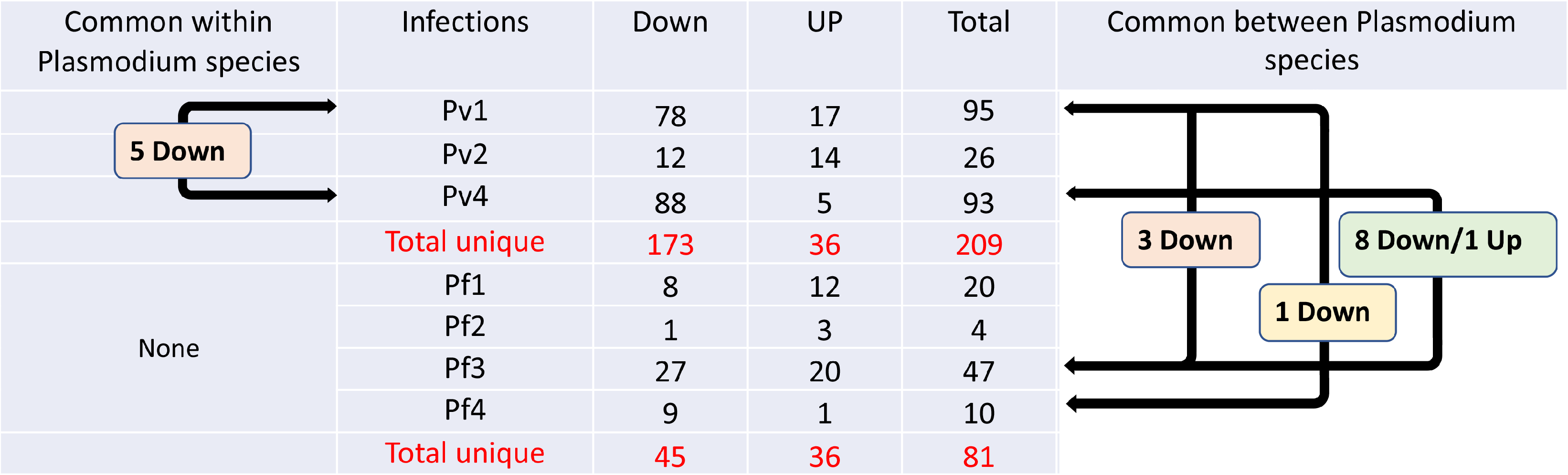
: Summary of *An. arabiensis* transcript numbers differentially expressed upon *P. vivax* and *P. falciparum* infection.

**Figure 1.**
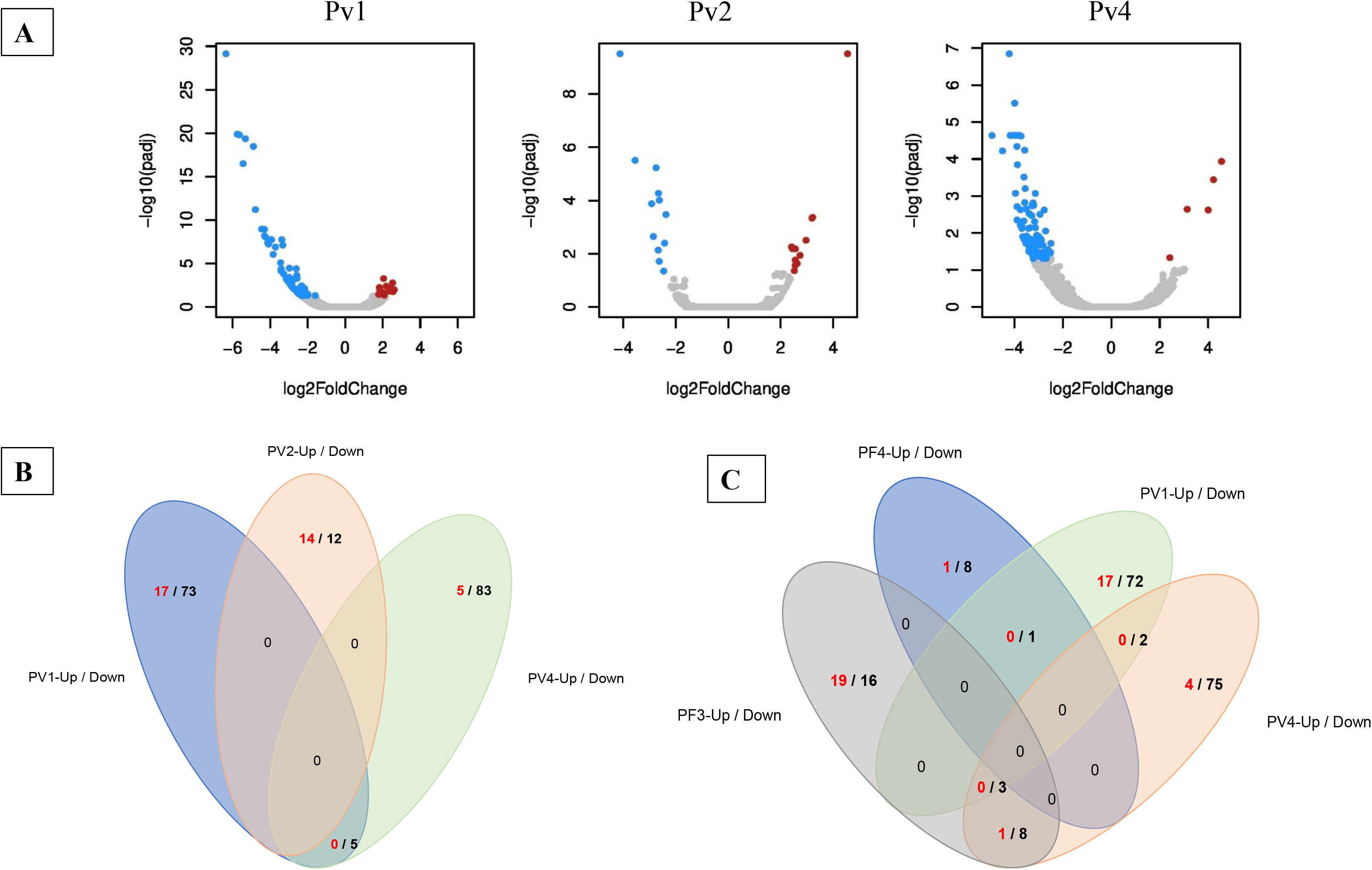
Volcano plots and numbers of *An. arabiensis* midgut differentially expressed genes upon *P. vivax* challenge. A) Plots of each pairwise comparison between infected and uninfected mosquito midgut samples. Transcripts were considered differentially expressed with a log2-fold change ≥ 1 and a corrected *p*-value (padj) < 0.05. Red dots display up-regulated genes and blue dots down-regulated genes. B and C) Venn diagrams highlighting shared up and down regulated mosquito genes among individual *P. vivax* infections (B), and among *P. vivax* and *P. falciparum* individual infections (C). Upregulated genes are labelled in red and downregulated ones in black.

### Differential gene expression in *An. arabiensis* midguts infected with *P. falciparum*

Eighty-one (81) transcripts were identified differentially regulated in *An. arabiensis* midguts in response to *P. falciparum*, 36 were up-regulated and 45 were down-regulated (Figure 2A,B and Table 1). Surprisingly, there was no common transcript between the four infections with *P. falciparum* isolates. The most down-regulated *An. arabiensis* gene was TRINITY_DN115268_c5_g1 in *Pf1* infection showing a log_2_FC at −4,154. This gene showed 34.11% identity to Membrane alanyl aminopeptidase (AMPM_HELVI) from *Heliothis virescens*. The second most down-regulated transcript, TRINITY_DN102799_c3_g1 in *Pf3* infection, had a log_2_FC at −3,624. This gene exhibits 69.24% identity to Cytochrome P450 from *Drosophila*. The most up-regulated transcript was TRINITY_DN114835_c4_g2 g, in *Pf3* infection with a log_2_FC at 4,113 and correspond to an unannotated gene.

**Figure 2.**
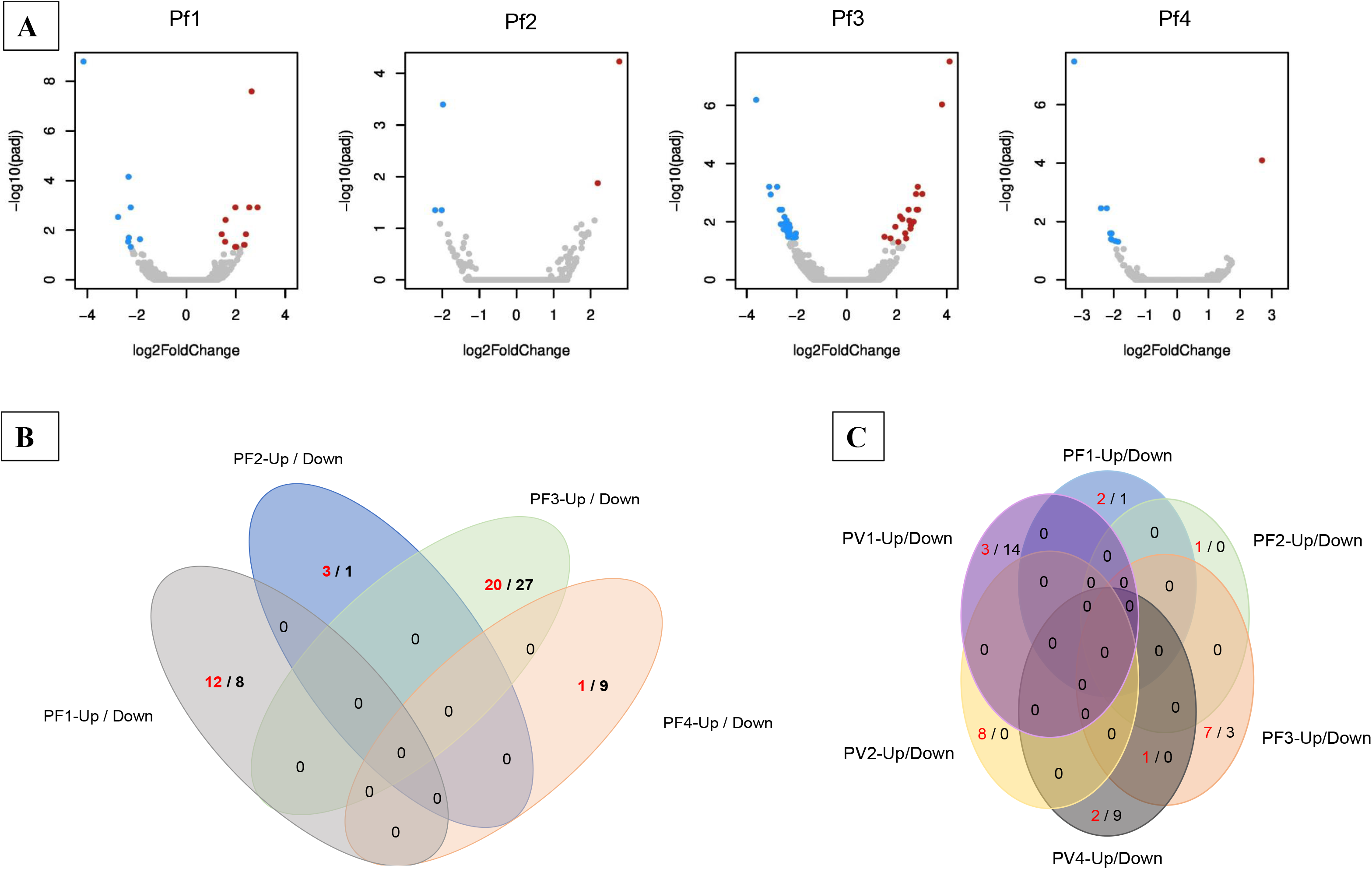
Volcano plots and numbers of *An. arabiensis* midgut differentially expressed genes upon *P. falciparum* challenge. A) Plots of each pairwise comparison between infected and uninfected mosquito midgut samples. Transcripts were considered differentially expressed with a log2-fold change ≥ 1 and a corrected *p*-value (padj) < 0.05. Red dots display up-regulated genes and blue dots down-regulated genes. B) Venn diagram highlighting shared up and down regulated mosquito genes among individual *P. falciparum* infections. C) Venn diagram highlighting shared up and down regulated mosquito lncRNAs among individual *P. vivax* and *P. falciparum* infections. Upregulated genes are in red and downregulated genes in black.

### Differential expression of long non-coding RNAs in *An. arabiensis* infected with *Plasmodium*

Among the 290 genes differentially expressed transcripts in response to either *P. vivax* or *P. falciparum*, 51 genes were classified as unannotated features. Using non-coding RNA identification programs, these 51 features were identified to be putative lncRNAs representing a total of 146 transcripts assembled by trinity. Among the 51 lncRNA genes, 14 were upregulated and 23 downregulated associated to *P. vivax* infection whereas 11 lncRNAs were upregulated and 4 downregulated associated to *P. falciparum* infection. No putative lncRNA was found in transcripts from the *Pf4* isolate. Among the upregulated lncRNAs, one was common to *An. arabiensis* infected with *Pv4* isolate and *Pf3* isolate. An overview of lncRNAs regulation is presented in Figure 2C and Table S5.

Of the 51 genes classified as unannotated features, 18 candidates were successfully mapped to the *An. arabiensis Dongola* genome at different loci. Three transcript features were identified as antisense lncRNAs (101849, 114154 and 106465) and are underlined in red in Table S5. They mapped respectively to Cuticular protein (TRINITY_DN101849_c1_g1 with five variants i1, i2, i3, i5 and i6 and showing log2FC at −4,25), Phosphoserine phosphatase (TRINITY_DN114154_c2_g1 with five variants i1 to i5 showing log2FC at −3.094) and Gambicin 1 (TRINITY_DN106465_c1_g3 with two variants i1 and i2 showing log2FC at −2.78). The 15 others were identified as sense lncRNA transcripts, some of them being associated to mosquito immune responsive genes. These included, Leucine-rich immune protein (Long) (TRINITY_DN112064_c0_g5_i1 with log2FC at −2.94), peptidoglycan recognition protein (TRINITY_DN107658_c0_g1_i1 with log2FC at −3.29), leucine-rich repeat protein (TRINITY_DN110940_c0_g1 with four variants i1, i5, i4 and i6 showing log2FC at 2.52) and Protein Kinase cGMP-dependent (TRINITY_DN113382_c1_g2 with four variants i1 to i4 showing log2FC at 2.38). A schematic alignment of the lncRNAs matching mosquito immune responsive genes is presented in Figure 3.

**Figure 3.**
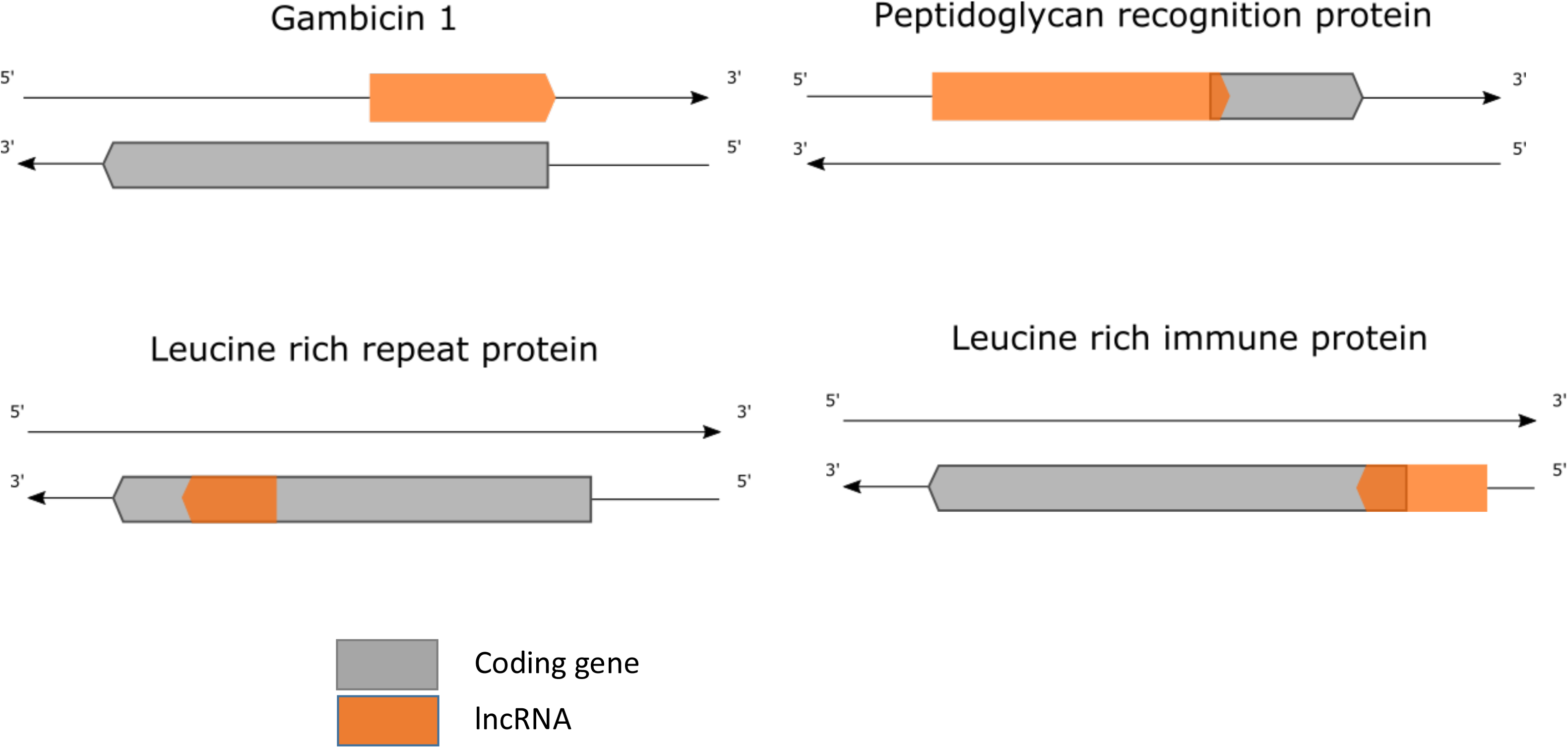
Overlapping of *An. arabiensis* regulated lncRNAs with known *Anopheles* immune responsive genes. ncRNA transcripts were aligned to *An. arabiensis* genes extracted from the Dongola AaraD1 genome and its annotation AaraD1.11.

### *An. arabiensis* midgut transcripts regulated by both *P. vivax* and *P. falciparum*

As presented above, there was a clear mosquito genotype to pathogen genotype interaction within parasite species. Our initial objective was to identify mosquito genes regulated similarly by both parasite as well as genes differently regulated by *P. vivax* and *P. falciparum*. We therefore looked in more details to *An. arabiensis* response to both human parasites, keeping in mind that the mosquito genotype (multiple F0 individuals) was not fixed. We identified 3 classes of transcripts: 1) those that were down regulated in both *P. vivax* and *P. falciparum* infected mosquitoes, 2) those that were upregulated in both groups, 3) those that were down regulated in *P. vivax* infected mosquitoes and upregulated in *P. falciparum* infected mosquitoes. No transcript upregulated in *P. vivax* infected mosquitoes and downregulated in *P. falciparum* infected mosquitoes was detected. The Venn diagram presented in Figure 1C highlighted the impact of parasite by mosquito genotype interaction for classes 1 and 2. Indeed, 3 transcripts were found down-regulated in both *Pv1* and *Pf3* infected mosquitoes, 8 transcripts down-regulated in *Pv4* and *Pf3* infected mosquitoes and an additional 1 down-regulated in *Pv1* and *Pf4,* whereas only one commonly up regulated transcript was shared between *Pv4 and Pf3*. From these data hierarchical clustering and heat maps were generated (Figure 4). In the heat maps, 11 class1 transcripts are globally downregulated in both *P. vivax* and *P. falciparum* infected mosquitoes (pink square) although data from two Pf4 midguts and one Pv2 midgut depart from the global trend. One transcript belonging to class 2 correspond to an unannotated gene (TRINITY_DN113404) further identified as a lncRNA (Table S5). Class 3 included three genes: Defensin1 (Def1), Peroxiredoxin-1,6 (PRDX6/PRX1), and Clavesin 1 (CLVS1). The overall differential regulation between *Pv* and *Pf* infected mosquitoes is nevertheless modest, with slight variation according to the parasite genotype. While Defensin1 and Peroxiredoxin have been associated to *Plasmodium*-mosquito interaction, the detection of a Clavesin related transcript is quite elusive in *Anopheles-Plasmodium* interaction as this gene family has been described as encoding Neuron-specific Lipid- and Clathrin-binding Sec14 Proteins (Katoh et al. 2009).

**Figure 4.**
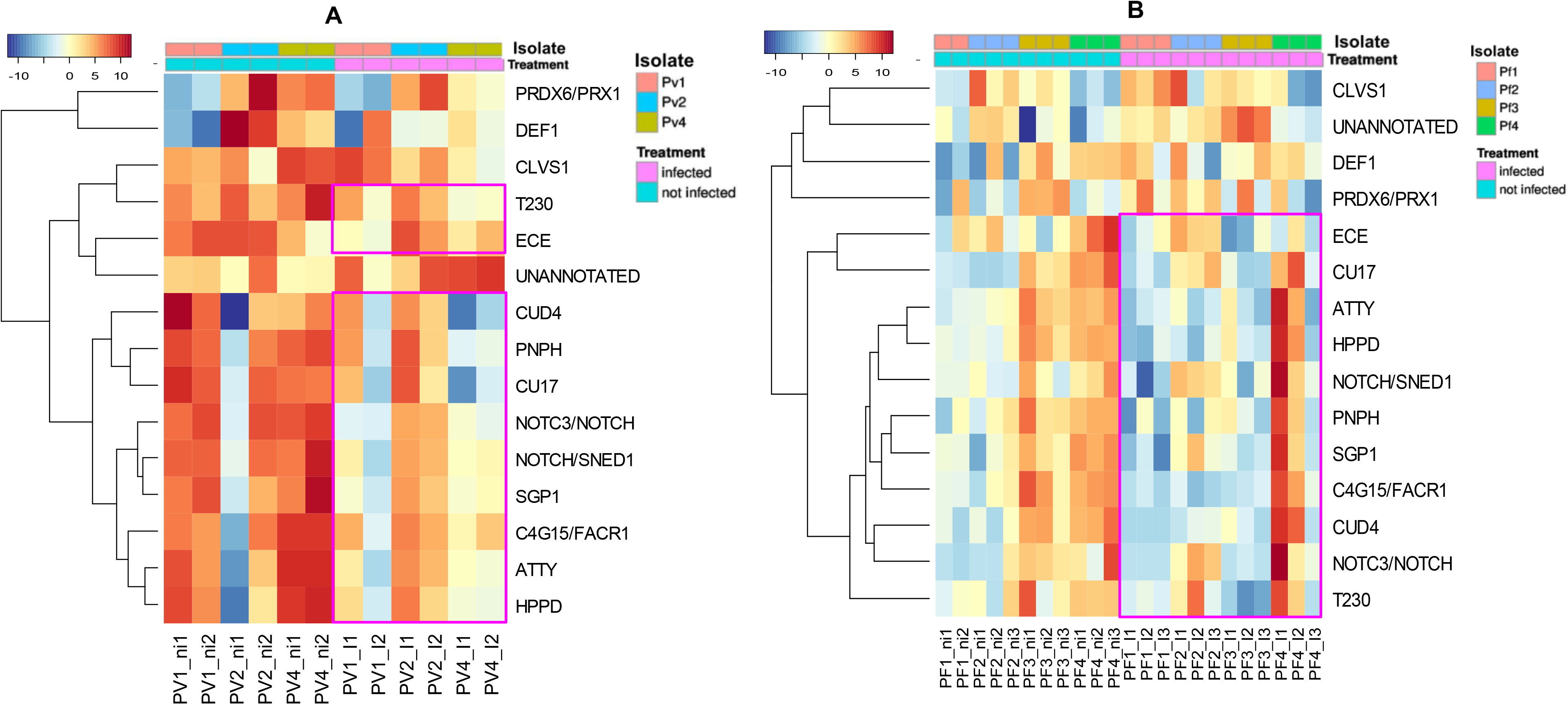
Heat map representation of shared differentially regulated genes in *An. arabiensis* mosquito midguts upon *Plasmodium* infection. A) *P. vivax* infection. B) *P. falciparum* infection. At the top of each heat map (A & B) pink color represents infected samples, while green color represents non infected samples (ie mosquitoes fed on inactivated parasites, see M&M). On panel A each *P. vivax* isolates (Pv1, Pv2 and Pv4) are color coded. Similarly, on panel B, each *P. falciparum* isolates (Pf1, Pf2, Pf3 and Pf4) are color coded. Each column is the measurement of change in gene expression (blue low expression and red high expression). Hierarchical clustering is represented by dendrogram trees. The pink rectangles delineate the 11 down-regulated genes in response to both *Pv* and *Pf* infection (T230: tryptophan 2,3-dioxygenase; ECE: Endothelin-converting enzyme; CUD4: Endocuticle structural glycoprotein ABD-4; PNPH: Purine nucleoside phosphorylase; CU17: Larval cuticle protein LCP-17; NOTC3/NOTCH: Neurogenic locus notch homolog protein 3; SGP1: Serine protease inhibitor I/II; NOTCH/SNED1: Neurogenic locus Notch protein; C4G15/FACR1: Cytochrome P450 4g15; HPPD: 4-hydroxyphenylpyruvate dioxygenase; ATTY: Tyrosine aminotransferase. The unannotated gene corresponds to an up-regulated lncRNA in response to both *P. vivax4* and *P. falciparum3* infection. Three genes were down-regulated in response to *Pv* and up-regulated in response to *Pf* (PRDX6/PRX1: Peroxiredoxin; CLVS1: Clavesin-1; DEF1: Defensin1).

### Gene Ontology classification of *An. arabiensis* genes regulated upon *Plasmodium* infection

Differentially expressed genes were further analyzed for functional classification using Gene Ontology (GO). GO enrichment analysis was performed using Trinity scripts and GOSeq 1.30.0 (Young et al. 2010) to provide a shortlist for potential genes of interest classified in three categories: biological process (BP), molecular function (MF) and cellular component (CC).

A summary of detailed annotations for each category corresponding to transcripts regulated in mosquito midgut upon each *Plasmodium* infection is depicted in Figure 5 and Table S6, using the highest number of enriched or depleted GO terms from *Pf3 and Pf1* (Figure 5A) *Pv1* and *Pv4* (Figures 5B-D).

**Figure 5.**
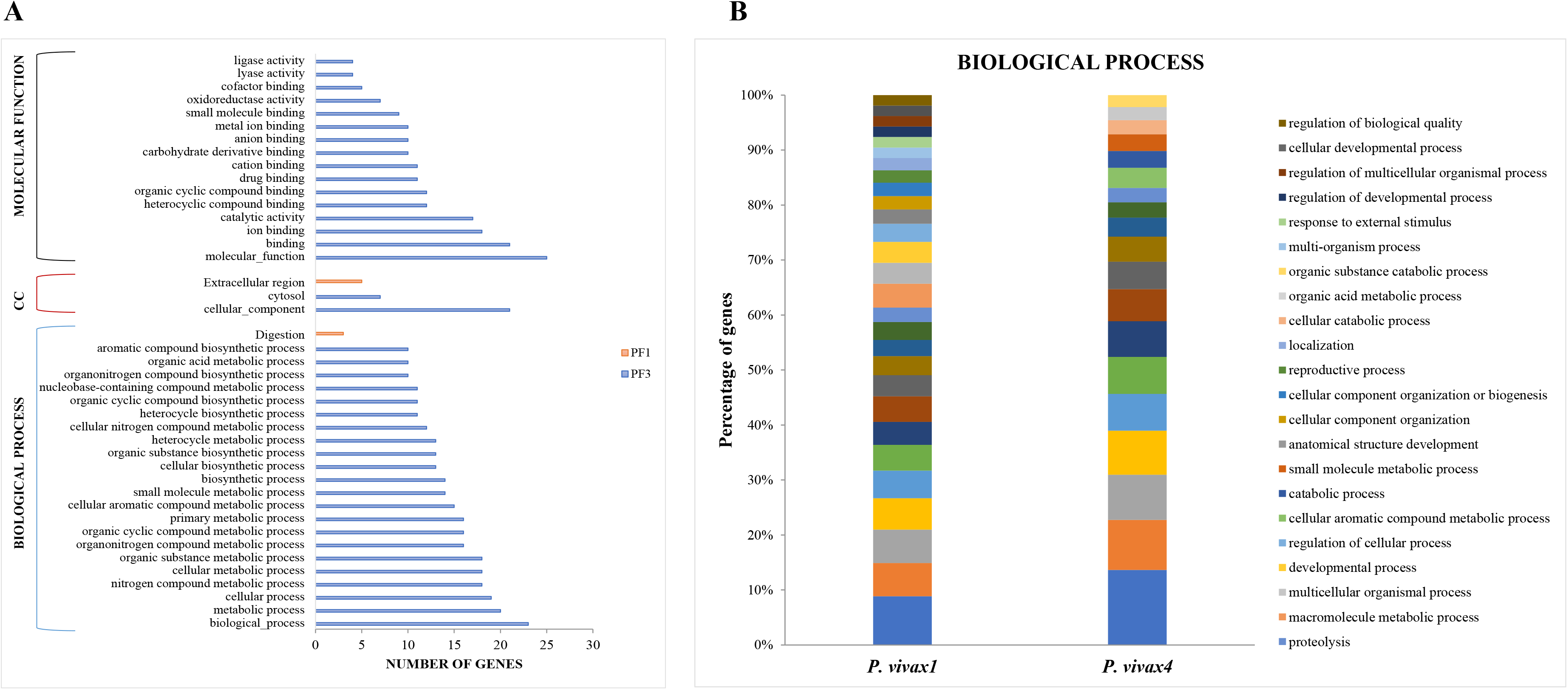
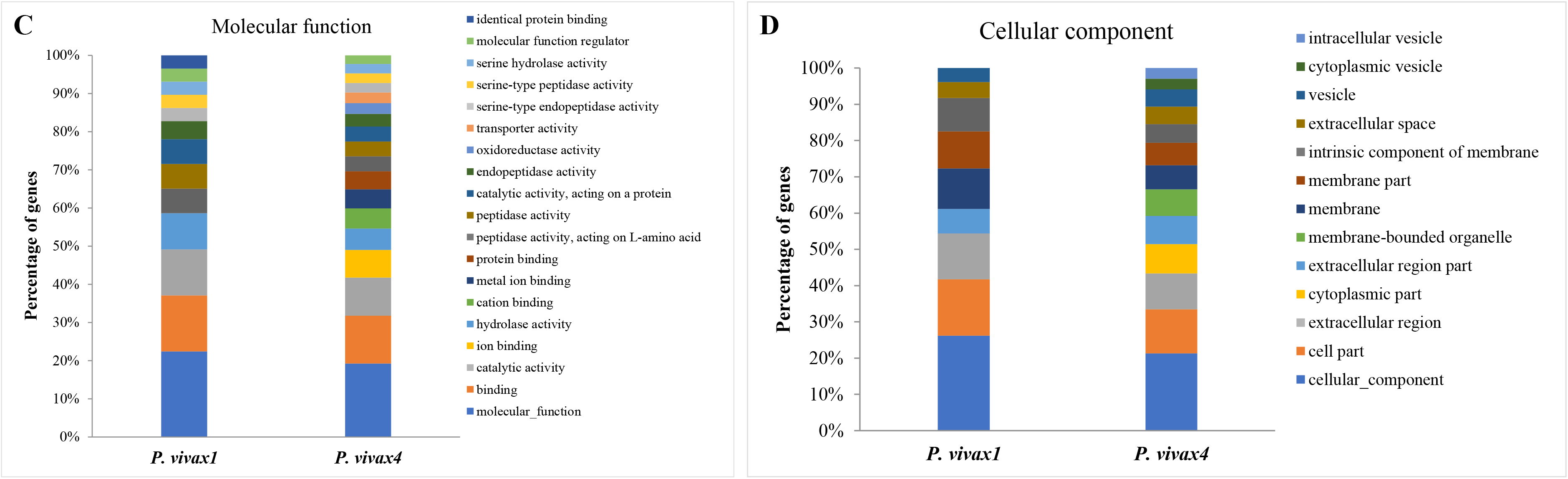
Gene ontology enrichment analysis of *An. arabiensis* genes 24 hours following *P. vivax and P. falciparum* infection. A false discovery rate of 0.05 (FDR< 0,05) was used for generating these graphs. A) Number of genes per category upon *P. falciparum* infection with isolates 1 and 3: biological process (BP), cellular component (CC) and molecular function (MF). B-D) Percentage of genes upon *P. vivax* infection with *Pv1* and *Pv4* isolates, according to biological process category (B), molecular function (C) and cellular component (D).

Metabolic and cellular processes were the most affected GO terms for biological process in mosquito midguts in response to *P. falciparum* and *P. vivax* infections: metabolic and biosynthetic processes for *P. falciparum* while proteolysis, multicellular organismal process, and developmental process for *P. vivax* (Figure 5A,B). The most changed GO terms for molecular function in mosquito midgut were binding activity in response to *P. falciparum* (Figure 5A) and mainly catalytic activity, peptidase and endopeptidase activity for *P. vivax* infections (Figure 5C). For the cellular component category, the enrichment analysis revealed the strongest enrichment of transcripts associated to cellular_component for both parasite species (Figures 5A and 5D).

## Discussion

We present in this report the first comparative investigation on *An. arabiensis* response to the two most important human malaria parasites *P. falciparum* and *P. vivax* using an RNA-Seq approach. Our study focused on RNAs differentially expressed in the mosquito midgut 24h post ingestion of infective *P. falciparum* and *P. vivax* gametocytes. This temporal and tissue specific window correspond to the migration of the parasite ookinetes across the mosquito midgut which correspond to the most vulnerable stage of the parasite both in numbers (Gouagna et al. 1998) and their accessibility for transmission blocking interventions (Saul 2007; Lavazec and Bourgouin 2008; Duffy 2021).

Working in a field station with natural populations of both *An. arabiensis* and *Plasmodium*, our RNA-Seq clearly demonstrates the importance of mosquito genotype to parasite genotype interaction in the building up of the mosquito transcriptional response. Indeed, each set of RNA-Seq data segregated according to the mosquito sample (Figure S3) except for *Pf1* and *Pf2* samples, that correspond to the same mosquito field population collected over the same week. Influence of parasite genotype is evidenced when looking at the number of differentially regulated transcripts in *Pf1* and *Pf2* infected mosquitoes, where *Pf1* led to 20 regulated transcripts between infected and uninfected mosquitoes whereas *Pf2* led to only 4 regulated transcripts (Table 1), despite similar gametocytemia in the two isolates (Table S1). Similarly, influence of parasite genotype can be seen within a *Plasmodium* species. Indeed, only 5 differentially regulated transcripts between infected and uninfected mosquitoes were common to the 3 *Pv* isolates, while none were found between the *Pf* isolates (Table 1).

Looking at the global response of *An. arabiensis* midgut tissue to *P. vivax* and *P. falciparum* infective gametocytes, our analysis reveals that wild *Plasmodium* isolates trigger the regulated expression of a limited number of transcripts, whether coding or not coding. Overall infection of *An. arabiensis* with *P. vivax* gametocytes regulate the expression of 209 transcripts while only 81 transcripts were regulated *by P. falciparum* gametocyte containing field isolates. The *P. falciparum* regulated transcript numbers are low compared to the higher numbers of regulated transcripts reported in transcriptomic studies of *An. gambiae* interaction with *P. falciparum* (Dong et al. 2006; Mead et al. 2012). However, these studies used established mosquito colonies (G3 or Keele strains) and referenced *P. falciparum* gametocyte producing strains (NF54 isolate or its 3D7 derivative clone). On the contrary, our data with *P. vivax* infected mosquitoes is in good agreement with results obtained with *A. aquasalis* (Santana et al. 2019) and *An. dirus* (Boonkaew et al. 2020), despite these studies used well established mosquito colonies. While the *An. dirus* study investigated whole mosquito transcriptome, revealing 313 regulated transcripts comparing mosquitoes fed on *P. vivax* gametocyte containing blood with mosquitoes fed on non -infected blood, the *An. aquasalis* study investigated mosquito midgut transcripts, revealing 49 regulated - transcripts comparing mosquitoes fed on *P. vivax* gametocyte containing blood with mosquitoes fed on inactivated gametocytes containing blood, similar to our experimental design. A globally limited number of midgut transcripts was also reported in *Aedes aegypti* upon challenge with Dengue virus (Raquin et al. 2017). Interestingly, or intriguingly, our results, in contrast to the *An. aquasalis* study, show that *P. vivax* down regulated a more significantly larger numbers of *An. arabiensis* transcripts than the number of up regulated ones. This could suggest that in a fully natural field combination *P. vivax* manipulates its host for the sake of its transmission.

Among the commonly down regulated genes, three genes belong to the catabolism of amino acid and notably tyrosine, which is known to be toxic in blood feeding arthropods when in excess as a result of the blood digestion (Sterkel et al. 2016). The observation that both *Plasmodium* species down regulated the expression of mosquito genes encoding amino acid degradation pathway is suggestive of parasite manipulation of the host for its own benefit. By contrast, it is surprising that both parasite species were down regulating a mosquito transcript encoding a purine nucleoside phosphorylase (PNP), involved in purine metabolism, owing that *Plasmodium* parasites are purine auxotrophic (in (Taylor et al. 2007). As intriguing is the recent report that *Caenorhabditis elegans* PNP ortholog is a negative regulator of the worm response to pathogens (Tecle et al. 2021). Two genes involved in mosquito immune response (encoding a P450 and a serpin) were as well down regulated. Such regulation is compatible with a pathogen evasion strategy to the mosquito immune response. Interestingly, *Plasmodium* parasites were also down regulating two genes encoding cuticular component (CU17 and a CUD4 family cuticular glycoprotein), in line with the needs for *Plasmodium* ookinetes to traverse the mosquito midgut peritrophic matrix before reaching the midgut epithelial cell (Sieber et al. 1991; Shahabuddin et al. 1994). Lastly is might seem odd that two neurogenic Notch encoding genes were down regulated in *Plasmodium* infected mosquito midgut. However, recent work revealed that the Notch-Delta signaling pathway is involved in *Ae. aegypti* response to DENV mosquito midgut infection (Serrato-Salas et al. 2018).

Our study led to the identification of several regulated transcripts corresponding to lncRNAs with only one common to *P. vivax* and *P. falciparum* infection. No match to annotated genes could be found for this upregulated lncRNA. Interestingly, among the lncRNAs four exhibit matches to immune responsive genes one being antisense to the *An. arabiensis* ortholog of *Gambicin 1*.

In conclusion, our comparative study *of P. vivax* and *P. falciparum* interaction with *An. arabiensis* is paving the way for a better understanding of the specific interaction of these malaria species with its anopheline vectors. Importantly, our study working with wild mosquitoes and wild parasites revealed a strong effect of mosquito by pathogen genotype interaction, which was not highlighted in any previous *Plasmodium-Anopheles* interaction studies. Our study highlights also for the first time, the regulation of lncRNAs, the function of which has yet to be determined. Additional functional analyses are required for selecting relevant genes as potential targets for *P. vivax and P. falciparum* transmission blocking approaches.

## Acknowledgments

We are grateful to volunteers from Andriba primary schools and their parents or guardians for participating in this study. We also address warm thanks to the medical teams of the Andriba and Antanimbary basic health center (CSB) for their assistance in collecting blood samples, notably Dr Richard and Odja, Andriba midwife. Many thanks to the whole local population who supported our study: Andriba city major, school directors, Funkuntany delegates, local military authorities, and health district authorities based in Maevatanana. A specific thanks to Jean-Pierre, guardian of the Andriba health center and his constant help for field mosquito larvae collections. We are also grateful to the technician staff from the Institut Pasteur de Madagascar: Tianasao from the malaria Unit, Manda, Fidelis and Miara from the medical entomology Unit, not forgetting our drivers and cooks, René, Bari and Hery.

## Author Contributions

Conceived the project and funding: CB. Designed the experiments: MTT, CB, M-AD. Provide local support: RG, MJ, AS, Performed the experiments: MTT, NP, CB, JMGY, SE, CP, ON. Analyzed the data: MTT, EK, CB. Contributed reagents/materials /analysis tools: CB. Draft the paper: MTT, EK, CB.

## Supporting files

**Table S1***Plasmodium* isolate characteristics and *An. arabiensis* infection rates

**Table S2**: RNA-Seq data summary: Statistics on sequenced reads before and after filtering by fasp and their kallisto pseudo-mapping percentage.

**Table S3.** Primer sequences.

**Table S4.** List of annotated and unannotated transcripts differentially expressed in *An. arabiensis* mosquito midgut in response to *P. vivax* and *P. falciparum* infection.

**Table S5**. List of *An. arabiensis* long non-coding RNAs differentially expressed in response to *P. vivax* and *P. falciparum* infection.

**Table S6.** Functional classification of *An. arabiensis* midgut transcripts using gene ontology analysis. The three categories are obtained upon *Pv* and *Pf* infection based on the FDR<0.05.

**Figure S1. Experimental design for mosquito infection by DMFA (Direct Membrane Feeding Assay) and RNA-Seq library production.** Three-to-five-day-old female mosquitoes were starved from sugar 16h prior blood-feeding on infective and non-infective blood containing *P. vivax* or *P. falciparum* gametocytes from field isolates. Individual mosquito midgut 24 post-blood-feeding were dissected in cold PBS and immediately transferred in Tri-Reagent. Samples were stored at −20°C for 3 to 4 days in the field station, before being placed at −80°C in Antananarivo and later fetched to Paris on dry ice. After total RNA extraction. cDNAs for RNA-Seq library were obtained from the selected RNAs and a total of 40 libraries submitted to sequencing.

**Figure S2.***P. vivax* **and***P. falciparum* **oocyst loads in infected***An. arabiensis*. The data represent the intensity of infection for each of the isolates used in the RNA-Seq analysis. Oocysts were detected on day 7 post infection. Additional details are presented in the M&M section and in Table S1. The left Y axis applies to all infections except Pf3 for which the right Y axis is drawn.

**Figure S3. Principal Components Analysis based on normalized counts from individual mosquito midguts infected with***P. vivax* **or***P. falciparum*. (A) PCA on global RNA-Seq patterns for mosquito fed on *P*. *vivax* infected and uninfected blood, PCA 1 and 2 displayed 58% of total variance; three clusters were clearly identified according to the *Pv* isolates. (B) PCA on global

RNA-Seq patterns for mosquito fed on *P*. *falciparum* infected and uninfected blood. Both PCA axis displayed 52% of total variance; three clusters were identified and PCA did not dissociate Pf1 and Pf2.

## References

Anstey, Nicholas M., Nicholas M. Douglas, Jeanne R. Poespoprodjo, and Ric N. Price. 2012. ‘Chapter Three - Plasmodium vivax: Clinical Spectrum, Risk Factors and Pathogenesis.’ in S. I. Hay, Ric Price and J. Kevin Baird (eds.), Advances in Parasitology (Academic Press).

Armistead, J. S., I. Morlais, D. K. Mathias, J. G. Jardim, J. Joy, A. Fridman, A. C. Finnefrock, A. Bagchi, M. Plebanski, D. G. Scorpio, T. S. Churcher, N. A. Borg, J. Sattabongkot, and R. R. Dinglasan. 2014. ‘Antibodies to a single, conserved epitope in Anopheles APN1 inhibit universal transmission of Plasmodium falciparum and Plasmodium vivax malaria, Infect Immun, 82: 818–29.

Ba, Hampâté, Craig W. Duffy, Ambroise D. Ahouidi, Yacine Boubou Deh, Mamadou Yero Diallo, Abderahmane Tandia, and David J. Conway. 2016. ‘Widespread distribution of Plasmodium vivax malaria in Mauritania on the interface of the Maghreb and West Africa’, Malaria Journal, 15: 80.

Baird, J. K. 2013. ‘Evidence and implications of mortality associated with acute Plasmodium vivax malaria’, Clin Microbiol Rev, 26: 36–57.

Battle, Katherine E., Tim C. D. Lucas, Michele Nguyen, Rosalind E. Howes, Anita K. Nandi, Katherine A. Twohig, Daniel A. Pfeffer, Ewan Cameron, Puja C. Rao, Daniel Casey, Harry S. Gibson, Jennifer A. Rozier, Ursula Dalrymple, Suzanne H. Keddie, Emma L. Collins, Joseph R. Harris, Carlos A. Guerra, Michael P. Thorn, Donal Bisanzio, Nancy Fullman, Chantal K. Huynh, Xie Kulikoff, Michael J. Kutz, Alan D. Lopez, Ali H. Mokdad, Mohsen Naghavi, Grant Nguyen, Katya Anne Shackelford, Theo Vos, Haidong Wang, Stephen S. Lim, Christopher J. L. Murray, Ric N. Price, J. Kevin Baird, David L. Smith, Samir Bhatt, Daniel J. Weiss, Simon I. Hay, and Peter W. Gething. 2019. ‘Mapping the global endemicity and clinical burden of Plasmodium vivax, 2000–17: a spatial and temporal modelling study’, The Lancet, 394: 332–43.

Boonkaew, Tippawan, Watcharakorn Mongkol, Sureerat Prasert, Pattaweeya Paochan, Saki Yoneda, Wang Nguitragool, Chalermpon Kumpitak, Jetsumon Sattabongkot, and Anchanee Kubera. 2020. ‘Transcriptome analysis of Anopheles dirus and Plasmodium vivax at ookinete and oocyst stages’, Acta Tropica, 207: 105502.

Boudin, C, A Diop, A Gaye, L Gadiaga, C Gouagna, I Safeukui, and S Bonnet. 2005. ‘Plasmodium falciparum transmission blocking immunity in three areas with perennial or seasonal endemicity and different levels of transmission’, Am J Trop Med Hyg, 73: 1090–95.

Bourgard, Catarina, Letusa Albrecht, Ana C. A. V. Kayano, Per Sunnerhagen, and Fabio T. M. Costa. 2018. ‘Plasmodium vivax Biology: Insights Provided by Genomics, Transcriptomics and Proteomics’, Frontiers in Cellular and Infection Microbiology, 8.

Bray, N. L., H. Pimentel, P. Melsted, and L. Pachter. 2016. ‘Erratum: Near-optimal probabilistic RNA-seq quantification’, Nat Biotechnol, 34: 888.

Bryant, D. M., K. Johnson, T. DiTommaso, T. Tickle, M. B. Couger, D. Payzin-Dogru, T. J. Lee, N. D. Leigh, T. H. Kuo, F. G. Davis, J. Bateman, S. Bryant, A. R. Guzikowski, S. L. Tsai, S. Coyne, W. W. Ye, R. M. Freeman, Jr., L. Peshkin, C. J. Tabin, A. Regev, B. J. Haas, and J. L. Whited. 2017. ‘A Tissue-Mapped Axolotl De Novo Transcriptome Enables Identification of Limb Regeneration Factors’, Cell Rep, 18: 762–76.

Camacho, C., G. Coulouris, V. Avagyan, N. Ma, J. Papadopoulos, K. Bealer, and T. L. Madden. 2009. ‘BLAST+: architecture and applications’, BMC Bioinformatics, 10: 421.

Chen, S., Y. Zhou, Y. Chen, and J. Gu. 2018. ‘fastp: an ultra-fast all-in-one FASTQ preprocessor’, Bioinformatics, 34: i884–i90.

Dinglasan, R. R., D. E. Kalume, S. M. Kanzok, A. K. Ghosh, O. Muratova, A. Pandey, and M. Jacobs-Lorena. 2007. ‘Disruption of Plasmodium falciparum development by antibodies against a conserved mosquito midgut antigen’, Proc Natl Acad Sci U S A, 104: 13461–6.

Dong, Y., R. Aguilar, Z. Xi, E. Warr, E. Mongin, and G. Dimopoulos. 2006. ‘Anopheles gambiae immune responses to human and rodent Plasmodium parasite species’, PLoS Pathog, 2: e52.

Duffy, Patrick E. 2021. ‘Transmission-Blocking Vaccines: Harnessing Herd Immunity for Malaria Elimination’, Expert Review of Vaccines: null-null.

Ewels, P., M. Magnusson, S. Lundin, and M. Kaller. 2016. ‘MultiQC: summarize analysis results for multiple tools and samples in a single report’, Bioinformatics, 32: 3047–8.

Fanello, C., F. Santolamazza, and A. della Torre. 2002. ‘Simultaneous identification of species and molecular forms of the Anopheles gambiae complex by PCR-RFLP’, Med Vet Entomol, 16: 461–4.

Finn, R. D., J. Clements, and S. R. Eddy. 2011. ‘HMMER web server: interactive sequence similarity searching’, Nucleic Acids Res, 39: W29–37.

Gouagna, L. C., B. Mulder, E. Noubissi, T. Tchuinkam, J. P. Verhave, and C. Boudin. 1998. ‘The early sporogonic cycle of Plasmodium falciparum in laboratory-infected Anopheles gambiae: an estimation of parasite efficacy’, Trop Med Int Health, 3: 21–8.

Goupeyou-Youmsi, Jessy, Tsiriniaina Rakotondranaivo, Nicolas Puchot, Ingrid Peterson, Romain Girod, Inès Vigan-Womas, Richard Paul, Mamadou Ousmane Ndiath, and Catherine Bourgouin. 2020. ‘Differential contribution of Anopheles coustani and Anopheles arabiensis to the transmission of Plasmodium falciparum and Plasmodium vivax in two neighbouring villages of Madagascar’, Parasites & Vectors, 13: 430.

Grabherr, M. G., B. J. Haas, M. Yassour, J. Z. Levin, D. A. Thompson, I. Amit, X. Adiconis, L. Fan, R. Raychowdhury, Q. Zeng, Z. Chen, E. Mauceli, N. Hacohen, A. Gnirke, N. Rhind, F. di Palma, B. W. Birren, C. Nusbaum, K. Lindblad-Toh, N. Friedman, and A. Regev. 2011. ‘Full-length transcriptome assembly from RNA-Seq data without a reference genome’, Nat Biotechnol, 29: 644–52.

Howes, R. E., K. E. Battle, K. N. Mendis, D. L. Smith, R. E. Cibulskis, J. K. Baird, and S. I. Hay. 2016. ‘Global Epidemiology of Plasmodium vivax’, Am J Trop Med Hyg, 95: 15–34.

Howes, R. E., S. A. Mioramalala, B. Ramiranirina, T. Franchard, A. J. Rakotorahalahy, D. Bisanzio, P. W. Gething, P. A. Zimmerman, and A. Ratsimbasoa. 2016. ‘Contemporary epidemiological overview of malaria in Madagascar: operational utility of reported routine case data for malaria control planning’, Malar J, 15: 502.

Isaacs, A. T., N. Jasinskiene, M. Tretiakov, I. Thiery, A. Zettor, C. Bourgouin, and A. A. James. 2012. ‘Transgenic Anopheles stephensi coexpressing single-chain antibodies resist Plasmodium falciparum development’, Proc Natl Acad Sci U S A, 109: E1922–30.

Katoh, Y., B. Ritter, T. Gaffry, F. Blondeau, S. Höning, and P. S. McPherson. 2009. ‘The clavesin family, neuron-specific lipid- and clathrin-binding Sec14 proteins regulating lysosomal morphology’, Journal of Biological Chemistry, 284: 27646–54.

Krogh, A., B. Larsson, G. von Heijne, and E. L. Sonnhammer. 2001. ‘Predicting transmembrane protein topology with a hidden Markov model: application to complete genomes’, J Mol Biol, 305: 567–80.

Kumari, Seena, Charu Chauhan, Sanjay Tevatiya, Deepak Singla, Tanwee Das De, Punita Sharma, Tina Thomas, Jyoti Rani, Deepali Savargaonkar, Kailash C. Pandey, Veena Pande, and Rajnikant Dixit. 2021. ‘Genetic changes of Plasmodium vivax tempers host tissue-specific responses in Anopheles stephensi’, Current Research in Immunology, 2: 12–22.

Lagesen, K., P. Hallin, E. A. Rødland, H. H. Staerfeldt, T. Rognes, and D. W. Ussery. 2007. ‘RNAmmer: consistent and rapid annotation of ribosomal RNA genes’, Nucleic Acids Res, 35: 3100–8.

Lavazec, C., C. Boudin, R. Lacroix, S. Bonnet, A. Diop, S. Thiberge, B. Boisson, R. Tahar, and C. Bourgouin. 2007. ‘Carboxypeptidases B of Anopheles gambiae as targets for a Plasmodium falciparum transmission-blocking vaccine’, Infect Immun, 75: 1635–42.

Lavazec, C., and C. Bourgouin. 2008. ‘Mosquito-based transmission blocking vaccines for interrupting Plasmodium development’, Microbes Infect, 10: 845–9.

Li, A., J. Zhang, and Z. Zhou. 2014. ‘PLEK: a tool for predicting long non-coding RNAs and messenger RNAs based on an improved k-mer scheme’, BMC Bioinformatics, 15: 311.

Love, M. I., W. Huber, and S. Anders. 2014. ‘Moderated estimation of fold change and dispersion for RNA-seq data with DESeq2’, Genome Biol, 15: 550.

Mathias, D. K., J. G. Jardim, L. A. Parish, J. S. Armistead, H. V. Trinh, C. Kumpitak, J. Sattabongkot, and R. R. Dinglasan. 2014. ‘Differential roles of an Anopheline midgut GPI-anchored protein in mediating Plasmodium falciparum and Plasmodium vivax ookinete invasion’, Infect Genet Evol.

Mead, E. A., M. Li, Z. Tu, and J. Zhu. 2012. ‘Translational regulation of Anopheles gambiae mRNAs in the midgut during Plasmodium falciparum infection’, BMC Genomics, 13: 366.

Menard, Didier, and Arjen Dondorp. 2017. ‘Antimalarial Drug Resistance: A Threat to Malaria Elimination’, Cold Spring Harbor Perspectives in Medicine, 7.

Mendes, AM, T Schlegelmilch, A Cohuet, P Awono-Ambene, M De Iorio, D Fontenille, I Morlais, GK Christophides, FC Kafatos, and D Vlachou. 2008. ‘Conserved mosquito/parasite interactions affect development of Plasmodium falciparum in Africa’, PLoS Pathog, 4: e1000069.

Miller, Louis H., Mary H. McGinniss, Paul V. Holland, and Paddy Sigmon. 1978. ‘The Duffy Blood Group Phenotype in American Blacks Infected with Plasmodium Vivax in Vietnam’, The American Journal of Tropical Medicine and Hygiene, 27: 1069–72.

Mukhtar, Maowia M., Omer A. Eisawi, Seth A. Amanfo, Elwaleed M. Elamin, Zeinab S. Imam, Faiza M. Osman, and Manasik E. Hamed. 2019. ‘Plasmodium vivax cerebral malaria in an adult patient in Sudan’, Malaria Journal, 18: 316.

Nguyen, Michele, Rosalind E. Howes, Tim C. D. Lucas, Katherine E. Battle, Ewan Cameron, Harry S. Gibson, Jennifer Rozier, Suzanne Keddie, Emma Collins, Rohan Arambepola, Su Yun Kang, Chantal Hendriks, Anita Nandi, Susan F. Rumisha, Samir Bhatt, Sedera A. Mioramalala, Mauricette Andriamananjara Nambinisoa, Fanjasoa Rakotomanana, Peter W. Gething, and Daniel J. Weiss. 2020. ‘Mapping malaria seasonality in Madagascar using health facility data’, BMC Medicine, 18: 26.

Niu, Guodong, Caio Franc̨a, Genwei Zhang, Wanlapa Roobsoong, Wang Nguitragool, Xiaohong Wang, Jetsumon Prachumsri, Noah S. Butler, and Jun Li. 2017. ‘The fibrinogen-like domain of FREP1 protein is a broad-spectrum malaria transmission-blocking vaccine antigen’, Journal of Biological Chemistry, 292: 11960–69.

Petersen, T. N., S. Brunak, G. von Heijne, and H. Nielsen. 2011. ‘SignalP 4.0: discriminating signal peptides from transmembrane regions’, Nat Methods, 8: 785–6.

Rabinovich, R. N., C. Drakeley, A. A. Djimde, B. F. Hall, S. I. Hay, J. Hemingway, D. C. Kaslow, A. Noor, F. Okumu, R. Steketee, M. Tanner, T. N. C. Wells, M. A. Whittaker, E. A. Winzeler, D. F. Wirth, K. Whitfield, and P. L. Alonso. 2017. ‘malERA: An updated research agenda for malaria elimination and eradication’, PLoS Med, 14: e1002456.

Rahimi, Bilal Ahmad, Ammarin Thakkinstian, Nicholas J. White, Chukiat Sirivichayakul, Arjen M. Dondorp, and Watcharee Chokejindachai. 2014. ‘Severe vivax malaria: a systematic review and meta-analysis of clinical studies since 1900’, Malaria Journal, 13: 481.

Ramasamy, M. S., R. Kulasekera, K. A. Srikrishnaraj, and R. Ramasamy. 1996. ‘Different effects of modulation of mosquito (Diptera:Culicidae) trypsin activity on the infectivity of two human malaria (Hemosporidia:Plasmodidae) parasites’, Journal of Medical Entomology, 33: 777–82.

Raquin, Vincent, Sarah Hélène Merkling, Valérie Gausson, Isabelle Moltini-Conclois, Lionel Frangeul, Hugo Varet, Marie-Agnès Dillies, Maria-Carla Saleh, and Louis Lambrechts. 2017. ‘Individual co-variation between viral RNA load and gene expression reveals novel host factors during early dengue virus infection of the Aedes aegypti midgut’, PLOS Neglected Tropical Diseases, 11: e0006152.

Santana, Rosa Amélia Gonçalves, Maurício Costa Oliveira, Iria Cabral, Rubens Celso Andrade Silva Junior, Débora Raysa Teixeira de Sousa, Lucas Ferreira, Marcus Vinícius Guimarães Lacerda, Wuelton Marcelo Monteiro, Patrícia Abrantes, Maria das Graças Vale Barbosa Guerra, and Henrique Silveira. 2019. ‘Anopheles aquasalis transcriptome reveals autophagic responses to Plasmodium vivax midgut invasion’, Parasites & Vectors, 12: 261.

Saul, A. 2007. ‘Mosquito stage, transmission blocking vaccines for malaria’, Curr Opin Infect Dis, 20: 476–81.

Serrato-Salas, J., J. Izquierdo-Sánchez, M. Argüello, R. Conde, A. Alvarado-Delgado, and H. Lanz-Mendoza. 2018. ‘Aedes aegypti antiviral adaptive response against DENV-2’, Dev Comp Immunol, 84: 28–36.

Service, M. W. 1993. Mosquito ecology: field sampling methods (Elsevier Applied Science: Elsevier: London; New York).

Shahabuddin, M., F. Lemos, M. Jacobs-Lorena, and D.C. Kaslow. 1994. “Mechanism of mosquito peritrophic matrix invasion by *Plasmodium* ookinetes and putative malaria transmission-blocking targets.” In 43rd Annual Meeting of the American Society of Tropical Medicine and Hygiene, 128. Cincinnati, Ohio, Nov13-17: Am. J. trop. Med. Hyg.

Sieber, K-P., M. Huber, D. Kaslow, S.M. Banks, M. Torii, M. Aikawa, and L.H. Miller. 1991. ‘The peritrophic membrane as a barrier: its penetration by *Plasmodium gallinaceum* and the effect of a monoclonal antibody to ookinetes’, Experimental Parasitology, 72: 145–56.

Simão, F. A., R. M. Waterhouse, P. Ioannidis, E. V. Kriventseva, and E. M. Zdobnov. 2015. ‘BUSCO: assessing genome assembly and annotation completeness with single-copy orthologs’, Bioinformatics, 31: 3210–2.

Smith, Ryan C, Joel Vega-Rodríguez, and Marcelo Jacobs-Lorena. 2014. ‘The Plasmodium bottleneck: malaria parasite losses in the mosquito vector’, Memórias do Instituto Oswaldo Cruz, 109: 644–61.

Solomon, Absra, Daniel Kahase, and Mihret Alemayehu. 2020. ‘Trend of malaria prevalence in Wolkite health center: an implication towards the elimination of malaria in Ethiopia by 2030’, Malaria Journal, 19: 112.

Soneson, C., M. I. Love, and M. D. Robinson. 2015. ‘Differential analyses for RNA-seq: transcript-level estimates improve gene-level inferences’, F1000Res, 4: 1521.

Sterkel, Marcos, Hugo D Perdomo, Melina G Guizzo, Ana Beatriz F Barletta, Rodrigo D Nunes, Felipe A Dias, Marcos H F. Sorgine, and Pedro L Oliveira. 2016. ‘Tyrosine Detoxification Is an Essential Trait in the Life History of Blood-Feeding Arthropods’, Current Biology, 26: 2188–93.

Taylor, E. A., A. Rinaldo-Matthis, L. Li, M. Ghanem, K. Z. Hazleton, M. B. Cassera, S. C. Almo, and V. L. Schramm. 2007. ‘Anopheles gambiae purine nucleoside phosphorylase: catalysis, structure, and inhibition’, Biochemistry, 46: 12405–15.

Tchuinkam, T., B. Mulder, K. Dechering, H. Stoffels, J.P. Verhave, M. Cot, P. Carnevale, J.H.E.T. Meuwissen, and V. Robert. 1993. ‘Experimental Infections of Anopheles-Gambiae with Plasmodium-Falciparum of Naturally Infected Gametocyte Carriers in Cameroon - Factors Influencing the Infectivity to Mosquitoes’, Tropical Medicine and Parasitology, 44: 271–76.

Tecle, Eillen, Crystal B. Chhan, Latisha Franklin, Ryan S. Underwood, Wendy Hanna-Rose, and Emily R. Troemel. 2021. ‘The purine nucleoside phosphorylase pnp-1 regulates epithelial cell resistance to infection in C. elegans’, PLOS Pathogens, 17: e1009350.

Wampfler, R., F. Mwingira, S. Javati, L. Robinson, I. Betuela, P. Siba, H. P. Beck, I. Mueller, and I. Felger. 2013. ‘Strategies for detection of Plasmodium species gametocytes’, PLoS ONE, 8: e76316.

Wang, L., H. J. Park, S. Dasari, S. Wang, J. P. Kocher, and W. Li. 2013. ‘CPAT: Coding-Potential Assessment Tool using an alignment-free logistic regression model’, Nucleic Acids Res, 41: e74.

Weiss, Daniel J., Tim C. D. Lucas, Michele Nguyen, Anita K. Nandi, Donal Bisanzio, Katherine E. Battle, Ewan Cameron, Katherine A. Twohig, Daniel A. Pfeffer, Jennifer A. Rozier, Harry S. Gibson, Puja C. Rao, Daniel Casey, Amelia Bertozzi-Villa, Emma L. Collins, Ursula Dalrymple, Naomi Gray, Joseph R. Harris, Rosalind E. Howes, Sun Yun Kang, Suzanne H. Keddie, Daniel May, Susan Rumisha, Michael P. Thorn, Ryan Barber, Nancy Fullman, Chantal K. Huynh, Xie Kulikoff, Michael J. Kutz, Alan D. Lopez, Ali H. Mokdad, Mohsen Naghavi, Grant Nguyen, Katya Anne Shackelford, Theo Vos, Haidong Wang, David L. Smith, Stephen S. Lim, Christopher J. L. Murray, Samir Bhatt, Simon I. Hay, and Peter W. Gething. 2019. ‘Mapping the global prevalence, incidence, and mortality of Plasmodium falciparum, 2000–17: a spatial and temporal modelling study’, The Lancet, 394: 322–31.

WHO. 2020. ‘World Malaria Report 2020’.

Young, M. D., M. J. Wakefield, G. K. Smyth, and A. Oshlack. 2010. ‘Gene ontology analysis for RNA-seq: accounting for selection bias’, Genome Biol, 11: R14.

Zhang, G., G. Niu, C. M. Franca, Y. Dong, X. Wang, N. S. Butler, G. Dimopoulos, and J. Li. 2015. ‘Anopheles Midgut FREP1 Mediates Plasmodium Invasion’, Journal of Biological Chemistry, 290: 16490–501.

