## Supplementary figures and images for "Differential transcriptomic response of *Anopheles arabiensis* to *Plasmodium vivax* and *Plasmodium falciparum* infection"

### Supplemental Figure 1

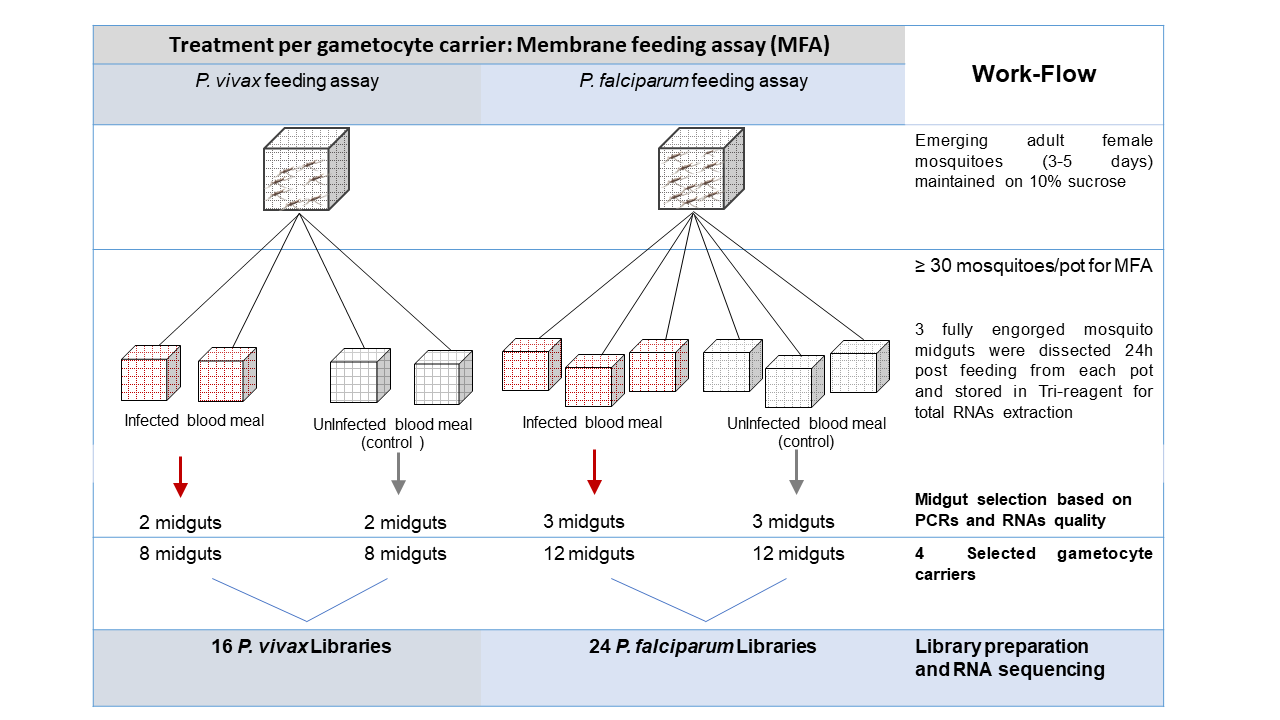

### Supplemental Figure 2

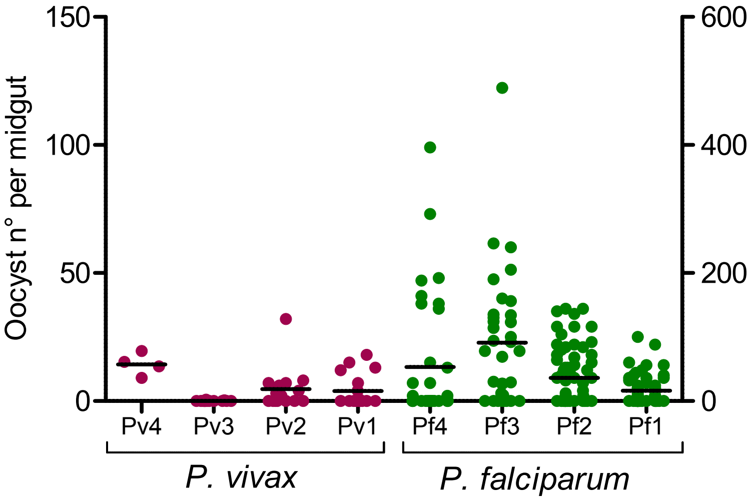

### Supplemental Figure 3

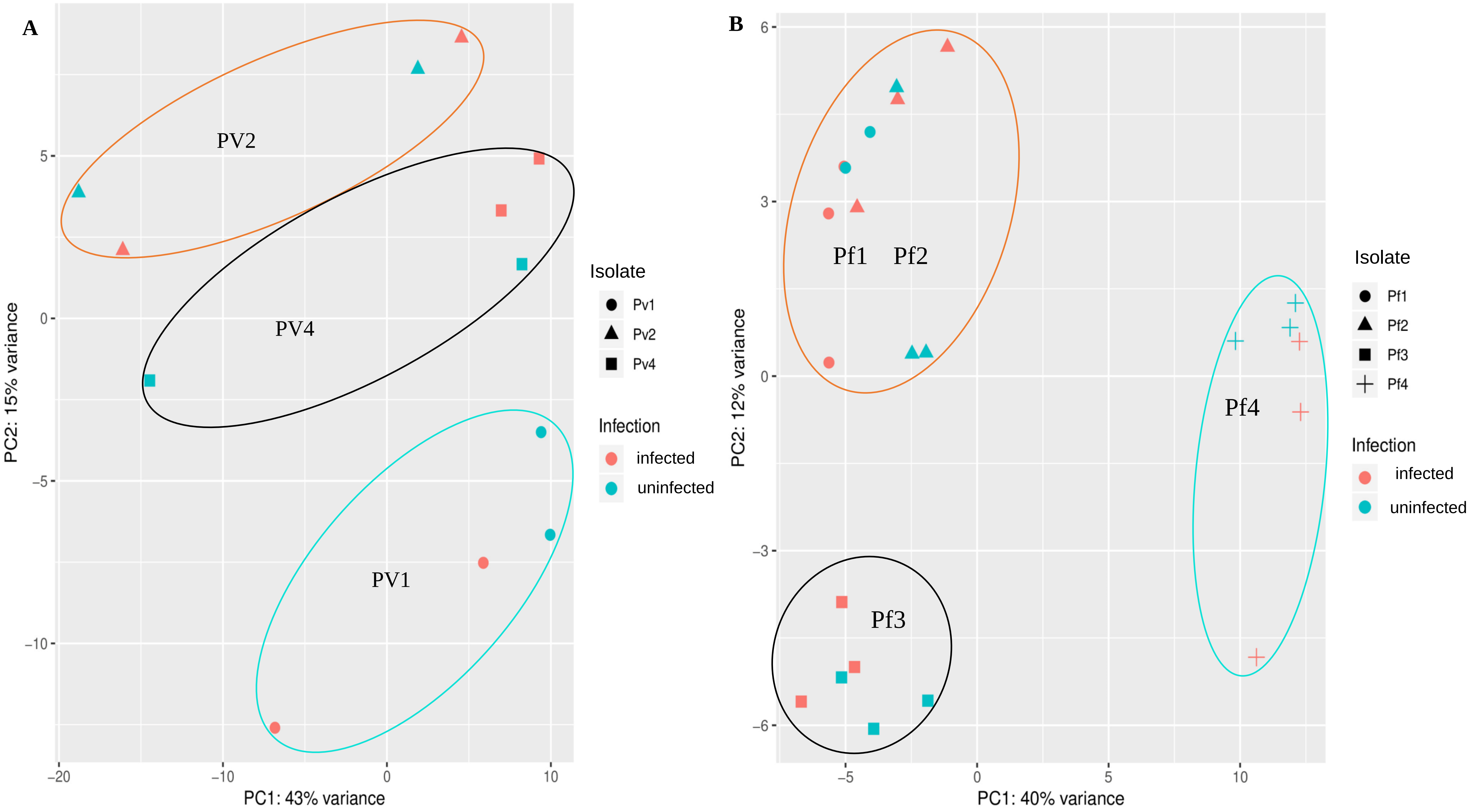
