## Supplemental Tables 1-3 for "Differential transcriptomic response of *Anopheles arabiensis* to *Plasmodium vivax* and *Plasmodium falciparum* infection"

**Table S1**. ***Plasmodium* isolate characteristics and *An. arabiensis* infection rates**

| **Isolate**  (Parasite) | **Asexual stage**  (µl) | **Sexual stage**  (µl) | **P-inf**  %(N/n) | **P-Ninf**  %(N/n) | **Oocyst**  **Mean** [range] |
| --- | --- | --- | --- | --- | --- |
| ***P. vivax*** |  |  |  |  |  |
| **Pv1** | 12639 | 315 | 38.9 (7/18) | 0 (0/6) | 10 [1-18] |
| **Pv2** | 2235 | 207 | 56.2 (9/16) | 0 (0/14) | 7.2 [2-32] |
| **Pv3** | 156 | 187 | 18.7 (3/16) | 0 (0/32) | 1.3 [1-2] |
| **Pv4** | 3333 | ND | 100 (4/4) | 0 (0/1) | 57.2 [36-78] |
| ***P. falciparum*** |  |  |  |  |  |
| **Pf1** | 255 | 271 | 61.6 (18/29) | 0 (0/32) | 6.6 [1-25] |
| **Pf2** | 32 | 300 | 75.8 (44/58) | 0 (0/21) | 14.3 [2-36] |
| **Pf3** | 11711 | 4118 | 76.5 (26/34) | 0 (0/6) | 119 [3-485] |
| **Pf4** | 0 | 1972 | 40 (14/35) | 0 (0/19) | 33.3 [2-99] |

Pv1 to Pv4 are the four selected *P. vivax* isolates, and Pf1 to Pf4 the four selected *P. falciparum* isolates used for mosquito infection and subsequent RNA-Seq investigation.

Asexual stages include ring, trophozoite and schizont stages while sexual stage corresponds to gametocytes. Stage numbers were estimated per blood µl from examination of Giemsa-stained thin blood smears on the day of mosquito infection.

P-inf and P-Ninf : prevalence of mosquito infection determined on day 7 post infection by detection of oocyst on mosquito midguts. P-inf: mosquitoes fed on “infective blood” ; P-Ninf mosquitoes fed on “non-infective blood”. The non-infective blood corresponds to heat-treated infective blood to inactivate gametocytes.

N : number of midguts harboring at least one oocyst ; n : the total number of observed mosquito midguts.

Oocyst mean: Mean number of oocysts from mosquitoes carrying at least one oocyst

ND: no gametocyte could be detected on the blood smear.

**Table S2. RNA-Seq total reads**

|  | **Before_filtering** | | | | **After_filtering** | | | | **kallisto** |
| --- | --- | --- | --- | --- | --- | --- | --- | --- | --- |
| **Index** | **gc_content** | **q30_rate** | **total_bases** | **total_reads** | **gc_content** | **q30_rate** | **total_bases** | **total_reads** | **percent_aligned** |
| Pf1-i1 | 0.53 | 0.95 | 6.38 | 58.02 | 0.52 | 0.96 | 5.39 | 56.8 | 91.18 |
| Pf1- i2 | 0.53 | 0.96 | 5.2 | 47.31 | 0.52 | 0.97 | 4.59 | 46.61 | 90.44 |
| Pf1- i3 | 0.53 | 0.96 | 6.87 | 62.43 | 0.52 | 0.97 | 5.98 | 61.31 | 91.84 |
| Pf1-ni1 | 0.53 | 0.96 | 4.91 | 44.63 | 0.52 | 0.97 | 4.24 | 43.97 | 58.99 |
| Pf1-ni2 | 0.52 | 0.97 | 6.78 | 61.66 | 0.51 | 0.97 | 6 | 60.93 | 89.35 |
| Pf1-ni3 | 0.52 | 0.95 | 7.99 | 72.64 | 0.51 | 0.96 | 6.9 | 70.38 | 90.22 |
| Pf2-i1 | 0.53 | 0.96 | 5.92 | 53.79 | 0.52 | 0.97 | 5.21 | 52.71 | 91.38 |
| Pf2-i2 | 0.53 | 0.96 | 6.56 | 59.62 | 0.52 | 0.97 | 5.65 | 58.69 | 92.32 |
| Pf2-i3 | 0.52 | 0.94 | 8.93 | 81.22 | 0.51 | 0.96 | 6.76 | 77.59 | 89.98 |
| Pf2-ni1 | 0.53 | 0.95 | 6.93 | 62.99 | 0.52 | 0.96 | 5.89 | 61.55 | 92.12 |
| Pf2-ni2 | 0.53 | 0.96 | 8.53 | 77.51 | 0.52 | 0.97 | 7.17 | 76.21 | 91.89 |
| Pf2-ni3 | 0.53 | 0.95 | 5.32 | 48.35 | 0.52 | 0.96 | 4.58 | 46.85 | 92.04 |
| Pf3-i1 | 0.53 | 0.97 | 3.48 | 31.62 | 0.52 | 0.97 | 3.08 | 31.3 | 91.77 |
| Pf3-i2 | 0.53 | 0.95 | 8.42 | 76.57 | 0.52 | 0.96 | 7.03 | 75.05 | 91.59 |
| Pf3-i3 | 0.53 | 0.96 | 5.89 | 53.54 | 0.52 | 0.97 | 5.03 | 52.56 | 90.6 |
| Pf3-ni1 | 0.53 | 0.96 | 7.74 | 70.37 | 0.52 | 0.97 | 6.65 | 69.33 | 88.98 |
| Pf3-ni2 | 0.53 | 0.94 | 8.36 | 75.99 | 0.52 | 0.96 | 6.78 | 72.94 | 91.12 |
| Pf3-ni3 | 0.53 | 0.96 | 5.27 | 47.87 | 0.52 | 0.97 | 4.55 | 47.19 | 90.97 |
| Pf4-i1 | 0.53 | 0.96 | 7.28 | 66.19 | 0.52 | 0.97 | 6.26 | 65.33 | 91.91 |
| Pf4-i2 | 0.53 | 0.95 | 7.39 | 67.22 | 0.52 | 0.97 | 6.01 | 65.19 | 92.35 |
| Pf4-i3 | 0.52 | 0.94 | 5.08 | 46.22 | 0.51 | 0.96 | 3.93 | 44.38 | 91.06 |
| Pf4-ni1 | 0.53 | 0.96 | 7.93 | 72.11 | 0.52 | 0.97 | 6.4 | 70.7 | 91.08 |
| Pf4-ni2 | 0.53 | 0.96 | 6.65 | 60.47 | 0.52 | 0.97 | 5.49 | 59.56 | 91.63 |
| Pf4-ni3 | 0.53 | 0.96 | 8.35 | 75.93 | 0.52 | 0.97 | 6.77 | 74.92 | 90.73 |
| Pv1-i1 | 0.53 | 0.96 | 7.39 | 67.17 | 0.52 | 0.97 | 6.59 | 66.06 | 90.98 |
| Pv1-i2 | 0.53 | 0.96 | 5.05 | 45.93 | 0.52 | 0.97 | 4.28 | 45.03 | 91.45 |
| Pv1-ni1 | 0.53 | 0.94 | 5.11 | 46.49 | 0.52 | 0.96 | 4.54 | 45.3 | 91.35 |
| Pv1-ni2 | 0.53 | 0.97 | 5.17 | 47.01 | 0.52 | 0.97 | 4.62 | 46.48 | 91.75 |
| Pv2-i1 | 0.53 | 0.94 | 9.45 | 85.95 | 0.52 | 0.96 | 8.06 | 83.08 | 90.72 |
| Pv2-i2 | 0.53 | 0.94 | 9.08 | 82.59 | 0.52 | 0.96 | 7.45 | 79.57 | 90.9 |
| Pv2-ni1 | 0.53 | 0.96 | 6.57 | 59.68 | 0.52 | 0.97 | 5.77 | 58.82 | 92.26 |
| Pv2-ni2 | 0.53 | 0.96 | 11.48 | 104.38 | 0.52 | 0.97 | 9.75 | 103.1 | 91.88 |
| Pv3-i1 | 0.53 | 0.97 | 6.72 | 61.06 | 0.52 | 0.98 | 5.89 | 60.43 | 90.45 |
| Pv3-i2 | 0.53 | 0.95 | 7.4 | 67.23 | 0.52 | 0.97 | 6.38 | 65.16 | 92 |
| Pv3-ni1 | 0.53 | 0.94 | 1.09 | 9.87 | 0.52 | 0.96 | 0.96 | 9.51 | 90.93 |
| Pv3-ni2 | 0.53 | 0.96 | 7.24 | 65.84 | 0.52 | 0.97 | 6.11 | 64.61 | 92.14 |
| Pv4-i1 | 0.53 | 0.95 | 6.11 | 55.59 | 0.52 | 0.96 | 5.33 | 54.34 | 93.34 |
| Pv4-i2 | 0.53 | 0.94 | 14.64 | 133.13 | 0.52 | 0.95 | 12.78 | 129.58 | 91.62 |
| Pv4-ni1 | 0.53 | 0.95 | 4.55 | 41.32 | 0.52 | 0.96 | 4.03 | 40.62 | 90.73 |
| Pv4-ni2 | 0.53 | 0.96 | 7.2 | 65.49 | 0.52 | 0.97 | 6.29 | 64.26 | 91.07 |

Index code :

The index are labelled as follows: Pf1: infection with P. falciparum isolate 1; Pf1-i1: RNA from midgut#1 fed on infective gametocytes ; Pf1-ni1: RNA from midgut#1 fed on non-infective gametocytes.

For *P. falciparum*, 4 isolates were used, producing each a set of 3 “infected” midguts and a set of 3 “Non-infected” midgut”.

For *P*. vivax: 4 isolates were used, producing each a set of 2 “infected” midguts and a set of 2 “Non-infected” midgut”.

Additional details are presented in the M&M section.

**Table S3. Primer sets used for mosquito identification by PCR, and gametocyte detection by RT-PCR**

| Target | Forward | Reverse |
| --- | --- | --- |
| An. gambiae rDNA -IGS | 5′-GTG TGC CCC TTC CTC GAT GT-3′ (**UN**) | 5′-CTG GTT TGG TCG GCA CGT TT-3′ (**GA**) |
| An. arabiensis rDNA -IGS | 5′-GTG TGC CCC TTC CTC GAT GT-3′ (**UN**) | 5′-AAG TGT CCT TCT CCA TCC TA-3′ (**AR**) |
| *Pf*s25 | 5′-GAA ATC CCG TTT CAT ACG CTT G-3′ | 5′-AGT TTT AAC AGG ATT GCT TGT ATC TAA-3′ |
| *Pv*s25 | 5′-ACA CTT GTG TGC TTG ATG TAT GTC-3′ | 5′-ACT TTG CCA ATA GCA CAT GAG CAA-3′ |

UN, AR and GA: Mosquito primers used to distinguish An. arabiensis from An. gambiae. UN: Universal common primer, AR: *A. arabiensis*-specific primer, GA : *A. gambiae*-specific primer (Fanello et al. 2002)

*Pf*s25 and *Pv*s25 : Primers designed for the specific amplification of the RNA encoding the major ookinete surface protein P25. (Wampfler, Mwingira et al. 2013)

References

Fanello, C., F. Santolamazza and A. della Torre (2002). "Simultaneous identification of species and molecular forms of the Anopheles gambiae complex by PCR-RFLP." Med Vet Entomol **16**(4): 461-464.

Wampfler, R., F. Mwingira, S. Javati, L. Robinson, I. Betuela, P. Siba, H. P. Beck, I. Mueller and I. Felger (2013). "Strategies for detection of Plasmodium species gametocytes." PLoS One **8**(9): e76316.
